# PomX, a ParA/MinD ATPase activating protein, is a triple regulator of cell division in *Myxococcus xanthus*

**DOI:** 10.1101/2020.12.14.422651

**Authors:** Dominik Schumacher, Andrea Harms, Silke Bergeler, Erwin Frey, Lotte Søgaard-Andersen

## Abstract

Cell division is precisely regulated to generate daughter cells of correct size and shape. In the social bacterium *Myxococcus xanthus*, the tripartite PomX/Y/Z complex directly stimulates positioning of the cytokinetic FtsZ-ring at midcell to mark the division site. The ∼15 MDa PomX/Y/Z complex associates with the nucleoid in a PomZ-dependent manner, translocates to midcell to stimulate FtsZ-ring formation, and undergoes fission during division. We demonstrate that PomX consists of two functionally distinct domains and has three functions. The N-terminal domain interacts with the ParA/MinD ATPase PomZ and stimulates PomZ ATPase activity. The C-terminal domain mediates PomX self-interaction, interaction to PomY, and serves as a scaffold for PomX/Y/Z complex formation. Moreover, the PomX/PomZ interaction is important for fission. These observations together with previous work support that the architecturally diverse ATPase activating proteins of ParA/MinD ATPases are highly modular and use the same mechanism to activate their cognate ATPase via a short positively charged N-terminal extension.

## Introduction

Accurate positioning of the cell division site ensures the formation of daughter cells of correct size and shape. In most bacteria, cell division initiates with positioning of the tubulin homolog FtsZ at the incipient division site^1^. Subsequently, FtsZ polymerizes to form a dynamic ring-like structure, the so-called (Fts)Z-ring, which serves as a scaffold for recruitment of other components of the division machinery^1^. Accordingly, systems that regulate division site positioning act at the level of Z-ring formation^2^. While the core components of the division machinery are conserved, the regulatory systems that ensure Z-ring positioning and, thus, the division site, are surprisingly diverse and also incompletely understood^2^. Interestingly, several of these systems have at their core a member of the ParA/MinD superfamily of P-loop ATPases that interacts with system-specific components to generate system-specific dynamic localization patterns that bring about correct Z-ring positioning^3^. These patterns include pole-to-pole oscillations in the *Escherichia coli* MinCDE system, bipolar gradient formation in the *Caulobacter crescentus* MipZ/ParB system, polar localization in the *Bacillus subtilis* MinCDJ/DivIVA system, and translocation across the nucleoid in the PomXYZ system of *Myxococcus xanthus*^4,5^.

ParA/MinD ATPases are key players in orchestrating the subcellular organization of bacterial cells and are not only involved in division site positioning but also in chromosome and plasmid segregation as well as positioning of other macromolecular complexes^3^. The function of ParA/MinD ATPases critically depends on the ATPase cycle during which they alternate between a monomeric form and a dimeric ATP-bound form^6-9^. Generally, ParA/MinD ATPases have a low intrinsic ATPase activity that is stimulated by a single cognate ATPase Activating Protein (AAP)^3^. While ParA/MinD ATPases share overall sequence conservation, the AAPs are much less conserved. The ParAB*S* systems for chromosome and plasmid segregation and the *E. coli* MinCDE system have provided fundamental insights into how ParA/MinD ATPase/AAP pairs interact to establish dynamic localization patterns: The AAP ParB stimulates ATPase activity of ATP-bound ParA dimers bound non-specifically to DNA^6,10,11^. In chromosome segregation systems, ParB binds to multiple *parS* sites at the origin of replication, forming a large complex^12-14^ while ATP-bound ParA dimers bind non-specifically to the nucleoid^6,9,11,15^. Upon replication, one of the duplicated ParB-*parS* complexes interacts with nucleoid-bound ParA dimers, thereby stimulating ATP hydrolysis and causing the release of ParA monomers from the nucleoid^10,16-18^. Released ParA monomers undergo nucleotide exchange and rebind to the nucleoid^10,16-18^. Repeated interactions between the large ParB-*parS* complex and nucleoid-bound ParA result in translocation of the ParB-*parS* complex to the opposite cell half^16,17,19,20^. In the MinCDE system, the AAP MinE, which is non-homologous to ParB, stimulates ATPase activity of membrane- and ATP-bound MinD dimers^21-25^. *In vivo* dimeric ATP-bound MinD forms a complex with the inhibitor of Z-ring formation MinC at the membrane^24,26-28^. Upon stimulation of MinD ATPase activity by MinE, the MinD/C complex is released from the membrane^21,22,29,30^. Subsequently, MinD undergoes nucleotide exchange and rebinds to the membrane together with MinC. These interactions result in the coupled pole-to-pole oscillations of the MinC/D complex and MinE^27,31^.

In the rod-shaped *M. xanthus* cells, Z-ring formation and positioning at midcell between two segregated chromosomes are stimulated by the tripartite PomX/Y/Z complex^32-34^. PomZ is a ParA/MinD ATPase while PomX and PomY separately have AAP activity and synergistically stimulate the low intrinsic ATPase activity of DNA- and ATP-bound dimeric PomZ^32,33^. PomX and PomY are non-homologous, and share homology with neither ParB nor MinE. The Pom proteins form a dynamically localized complex with an estimated size of ∼15 MDa *in vivo* that is visible as a cluster by epifluorescence microscopy^32^. This complex associates with the nucleoid via PomZ, and early during the cell cycle, it is positioned on the nucleoid away from midcell, i.e. off-center, close to the new cell pole^32^. Subsequently, the complex translocates by biased random motion on the nucleoid to the midnucleoid, which coincides with midcell. At midnucleoid, the PomXYZ complex undergoes constrained motion and stimulates Z-ring formation. Intriguingly, during cell division, the PomX/Y/Z complex undergoes fission, with the two “portions” segregating to the two daughters^32^.

The Pom proteins interact in all three pairwise combinations *in vitro* and PomX is essential for cluster formation by PomY and PomZ *in vivo*^32^: *In vitro* PomX spontaneously polymerizes to form filaments, and alone forms a cluster *in vivo*. PomY bundles PomX filaments *in vitro*; *in vivo* PomY is recruited by PomX to form a PomX/Y complex, which is not associated with the nucleoid and stalled somewhere in cells. ATP- and DNA-bound PomZ dimers, but not monomeric PomZ, are recruited to the PomX/Y complex by interactions with PomX as well as PomY, resulting in the association of the PomX/Y/Z complex to the nucleoid. Due to the AAP activity of PomX and PomY, PomZ is rapidly turned over in the PomX/Y/Z complex and released in a monomeric form to the cytosol. Motion of the PomX/Y/Z complex depends on non-specific DNA binding and ATP hydrolysis by PomZ^32^. Translocation to midnucleoid and constrained motion at midnucleoid arise from the continuous turnover of PomZ in the complex together with the diffusive PomZ flux on the nucleoid into the PomX/Y/Z complex^32^. Finally, while all three Pom proteins are important for Z-ring formation at midcell, PomY and PomZ in the PomX/Y/Z complex are thought to be directly involved in recruiting FtsZ to the division site based on protein-protein interaction analyses^32^.

The Pom system displays unusual spatiotemporal dynamics and is also unusual because it incorporates two AAPs. It remains elusive how PomX and PomY stimulate PomZ ATPase activity. Here, we address the function of PomX *in vivo* and *in vitro* and show that it is a triple regulator of cell division with the AAP activity residing in the N-terminal domain, the C-terminal domain serving as a scaffold for PomX/Y/Z complex formation, and the PomX/Z interaction being important for PomX/Y/Z complex fission during division. Moreover, our findings support the notion that that AAPs of ParA/MinD ATPases use the same mechanism to activate their cognate ATPase.

## Results

### PomX consists of two functional domains with distinct activities

PomX and PomY co-occur with PomZ in Cystobacterineae^32^. PomZ sequences are highly conserved, while PomX and PomY sequences are more divergent (Fig. 1a). In *M. xanthus* PomX, the region from residues 27-182 is Ala and Pro-rich (23% Pro; 19% Ala) and the region from residues 222-401 contains a coiled-coil domain (Fig. 1b). Although PomX homologs are of different length, their overall architecture is similar and with a high level of similarity in the C-terminal regions while the N-terminal regions vary in length and similarity (Fig. 1b). To analyze how PomX functions, we divided PomX into two parts. From hereon, we refer to these two parts as the N-terminal domain (PomX^N^, residues 1-213) and the C-terminal domain (PomX^C^, residues 214-404) (Fig. 1c).

**Fig. 1.**
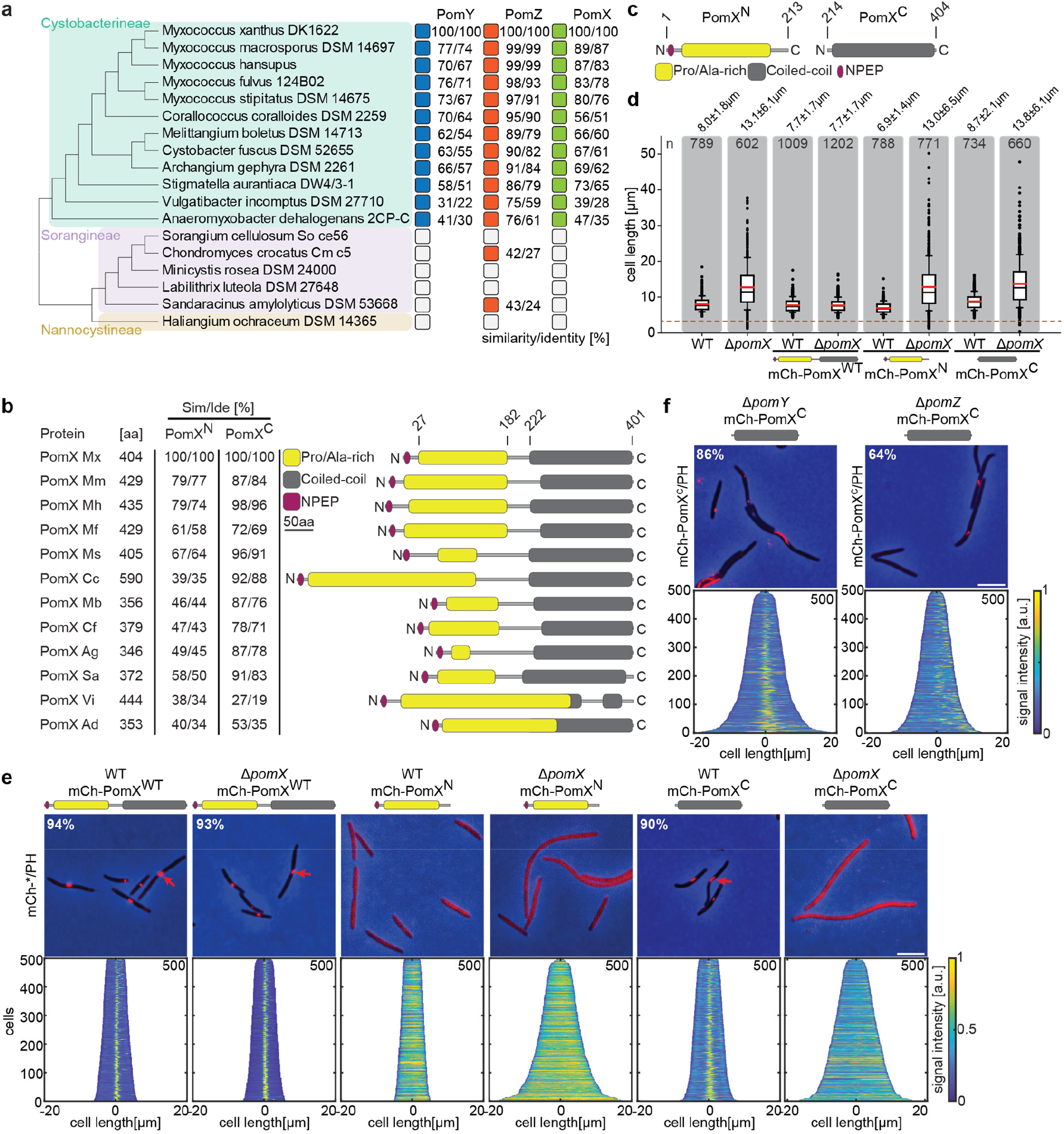
PomX consists of two domains that are both required for function. **a** Similarity and identity analysis of PomX, PomY, and PomZ homologs. The three Myxococcales suborders are indicated. An open box indicates that a homolog is not present. **b** Similarity and identity of PomX domains in different PomX homologs. Similarity and identity were calculated based on the domains of *M. xanthus* PomX shown in c. **c** PomX truncations used in this study. Numbers on top indicate the start and stop positions of the truncations relative to full-length PomX^WT^. **d** Cell length distribution of cells of indicated genotypes. Cells below stippled line are minicells. Numbers indicate mean cell length±STDEV. In the boxplots, boxes include the 25^th^ and the 75^th^ percentile, whiskers data points between the 10% and 90% percentile, outliers are shown as black dots. Black and red lines indicate the median and mean, respectively. Number of analyzed cells is indicated. In the complementation strains, *pomX* alleles were expressed from plasmids integrated in a single copy at the *attB* site. **e** Fluorescence microscopy of cells of indicated genotypes. Phase-contrast and fluorescence images of representative cells were overlayed. Numbers indicate fraction of cells with fluorescent cluster. Demographs show fluorescence signals of analyzed cells sorted according to length and with off-center signals to the right. Numbers in upper right indicate number of cells used to create demographs. Scale bar, 5µm. **f** Fluorescence microscopy of cells of indicated genotypes. Images of representative cells and demographs were created as in **e**. Scale bar, 5µm. For experiments in **d, e** and **f** similar results were obtained in two independent experiments.

We fused the two PomX domains to mCherry (mCh) and expressed them ectopically (Supplementary Fig. 1a). mCh-PomX^N^, mCh-PomX^C^, and full-length mCh-PomX^WT^ accumulated at or above PomX^WT^ levels in Δ*pomX* and *pomX*^+^ cells (Supplementary Fig. 1b). WT *M. xanthus* cells have a cell length of 8.0±1.8µm (mean±standard deviation (STDEV)), while Δ*pomX* cells are filamentous with a length of 13.1±6.1µm and also generate DNA-free minicells (Fig. 1d). As previously reported^32^, mCh-PomX^WT^ complemented the division defect of the Δ*pomX* mutant while the truncated variants did not.

mCh-PomX^WT^ formed a single well-defined cluster in 93% and 94% of Δ*pomX* and WT cells, respectively (Fig. 1e). These clusters localized in the off-center position (defined as clusters outside the midcell region at 50±5% of cell length) in short cells and at midcell in long cells (Fig. 1e). The truncated mCh-PomX variants displayed diffuse localization in Δ*pomX* cells (Fig. 1e). However, mCh-PomX^C^ but not mCh-PomX^N^ formed a single cluster in 90% of *pomX*^+^ cells and localized as mCh-PomX^WT^ (Fig. 1e). This cluster formation by mCh-PomX^C^ was independent of PomY and PomZ (Fig. 1f; Supplementary Fig. 1b), supporting that mCh-PomX^C^ is integrated into the PomX/Y/Z cluster via interaction with PomX. As described ^32^, the PomX clusters were more elongated in the absence of PomY (Fig. 1f). The incorporation of PomX^C^ into the PomX/Y/Z complex interfered with neither PomX/Y/Z complex formation nor function (Fig. 1d, e). We conclude that both PomX domains are essential for function and that mCh-PomX^C^ can integrate into the PomX/Y/Z complex via interaction with PomX while mCh-PomX^N^ cannot.

### PomX AAP activity resides in PomX^N^

We used the bacterial adenylate cyclase two-hybrid system (BACTH) to test for protein-protein interactions involving the two PomX domains. In agreement with previous observations using purified proteins, full-length PomX^WT^ self-interacted and interacted with PomY (Fig. 2a). PomX^C^ self-interacted and also interacted with full-length PomX and PomY, while PomX^N^ neither self-interacted nor interacted with PomX^WT^, PomX^C^, or PomY. To test for interactions with PomZ, we used two PomZ variants: PomZ^WT^ and PomZ^D90A^, which is locked in the DNA-binding, dimeric, ATP-bound form that interacts strongly with the PomX/PomY cluster *in vivo*^32^. PomX^WT^ and PomX^N^ interacted with PomZ^WT^ and PomZ^D90A^ while PomX^C^ did not; also, PomZ^D90A^ interacted more strongly with PomX^N^ than PomZ^WT^ (Fig. 2a).

**Fig. 2.**
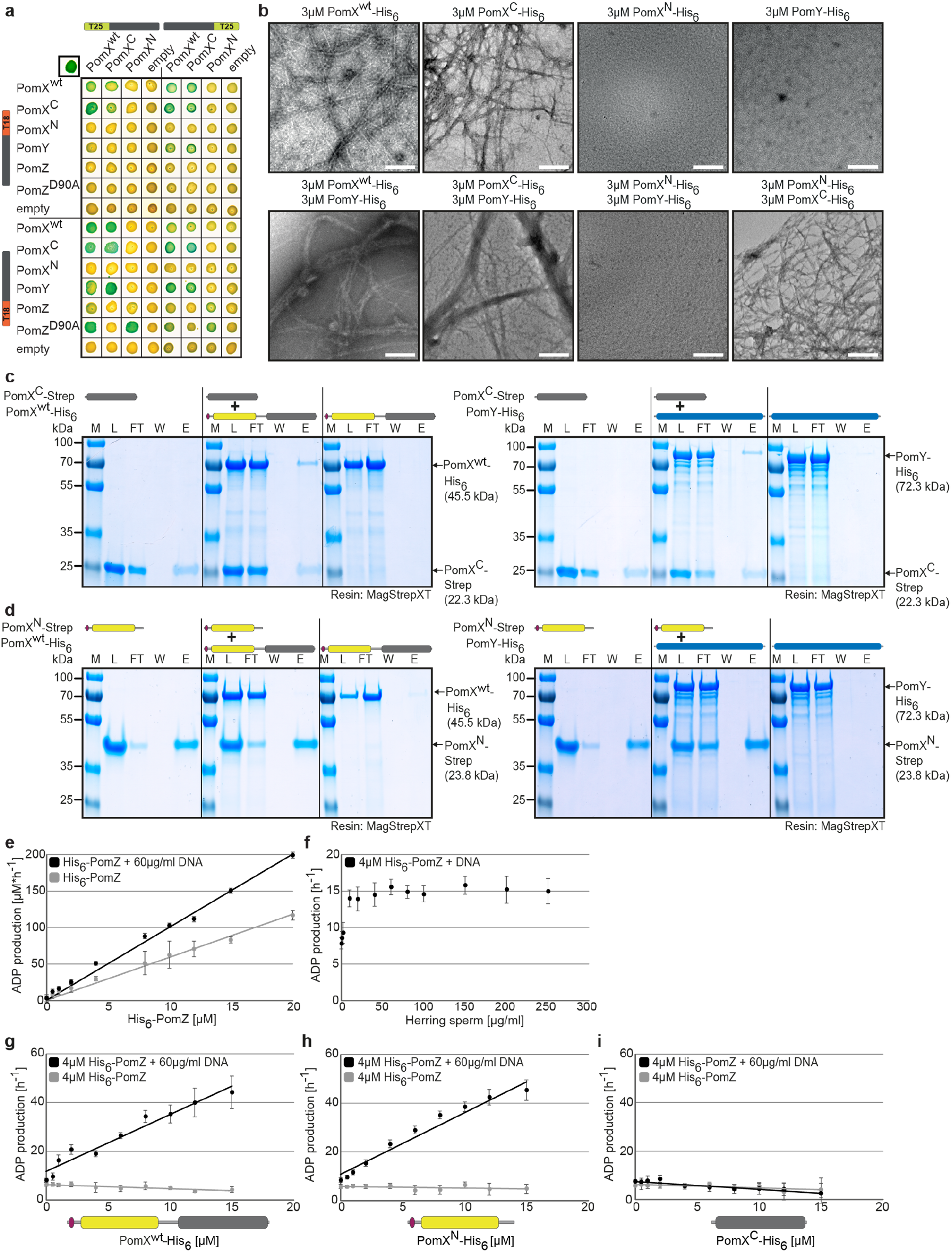
PomX^C^ interacts with PomX and PomY while PomX^N^ stimulates PomZ ATPase activity. **a** BACTH analysis of interactions between Pom proteins. The indicated protein fragments were fused to T18 and T25 as indicated. Blue colony indicates an interaction, white no interaction. Positive control in upper left corner, leucine zipper of GCN4 fused to T25 and T18. For negative controls, co-transformations with empty plasmids were performed. Images show representative results and were performed in three independent experiments. **b** TEM images of negatively stained purified proteins. Proteins were applied to the EM grids alone or after mixing in a 1:1 molar ratio as indicated before staining. Scale bar, 200nm. Images show representative results of several independent experiments. **c, d** *In vitro* pull-down experiments with purified PomX^C^-Strep, PomX^N^-Strep, PomX^WT^-His_6_, and PomY-His_6_. Instant Blue™-stained SDS-PAGE shows load (L), flow-through (FL), wash (W), and elution (E) fractions using MagStrep XT beads in pull-down experiments with 10µM of indicated proteins alone or pre-mixed as indicated on top. Molecular size markers are shown on the left and proteins analysed on the right together with their calculated MW. Note that PomX^WT^-His_6_ ^32^ and PomX^N^-Strep migrate aberrantly and according to a higher MW. All samples in a panel were analysed on the same gel and black lines are included for clarity. Experiments were repeated in two independent experiments with similar results. **e-i** His_6_-PomZ ATPase activity. ADP production rate was determined in an NADH-coupled photometric microplate assay in the presence of 1 mM ATP at 32°C. DNA and PomX variants were added as indicated. Spontaneous ATP hydrolysis and NADH consumption was accounted for by subtracting the measurements in the absence of His_6_-PomZ. Data points show the mean±STDEV calculated from six independent measurements.

To test for interactions between the Pom proteins *in vitro*, we purified tagged variants of PomX^WT^, PomX^N^, PomX^C^, PomY, and PomZ (Supplementary Fig. 2a). We confirmed by negative stain transmission electron microscopy (TEM) that PomX^WT^-His_6_ formed filaments that were bundled by PomY-His_6_, while PomY-His_6_ on its own did not form higher-order structures (Fig. 2b)^32^. Consistently, when analysed separately in high-speed centrifugation experiments, 93-95% of PomX^WT^-His_6_ and 29-35% of PomY-His_6_ were recovered in the pellet fraction, while 71-86% and 48%-49%, respectively of PomX^WT^-His_6_ and PomY-His_6_ were in the pellet fraction when mixed in equimolar amounts (Supplementary Fig. 2b, c, d).

As noted for PomX^WT32^, PomX^N^-His_6_ (molecular weight (MW) of monomer: 24.3 kDa) also migrated aberrantly in SDS-PAGE (Supplementary Fig. 2a). By size exclusion chromatography (SEC), the majority of PomX^N^-His_6_ eluted corresponding to a globular protein with a MW of ∼136 kDa and in a smaller peak corresponding to a MW of ∼306 kDa (Supplementary Fig. 2e). PomX^N^-His_6_ neither formed higher-order structures by TEM nor in high-speed centrifugation experiments (Fig. 2b; Supplementary Fig. 2c, d). Because PomX^N^ does not self-interact in BACTH, migrates aberrantly by SDS-PAGE, and is rich in Ala/Pro residues and, therefore, may not have a globular conformation, it is unclear whether SEC reflects the formation of PomX^N^ oligomers. PomX^C^-His_6_ in SDS-PAGE migrated at the expected size (MW of monomer: 22.8 kDa) (Supplementary Fig. S2a); however, the protein could not be analyzed by SEC because it did not enter the matrix. Accordingly, PomX^C^-His_6_ spontaneously formed filaments visible by TEM and mostly accumulated in the pellet fraction in high-speed centrifugation experiments (Fig. 2b; Supplementary Fig. 2b, d). PomY-His_6_ bundled the PomX^C^-His_6_ filaments and was enriched in the pellet fraction in centrifugation assays in the presence of PomX^C^-His_6_ (Fig. 2b; Supplementary Fig. 2b). Interactions between PomX^N^-His_6_ and PomX^WT^-His_6_, PomX^C^-His_6_ and PomY-His_6_ were observed by neither TEM nor high-speed centrifugation (Fig. 2b; Supplementary Fig. 2c, d). Finally, we confirmed in pull-down experiments using truncated PomX-Strep variants (Supplementary Fig. 2a) that PomX^WT^-His_6_ and PomY-His_6_ interact with PomX^C^-Strep (Fig. 2c) but not with PomX^N^-Strep (Fig. 2d) and that PomX^C^-His_6_ did not interact with PomX^N^-Strep (Supplementary Fig. 2f).

To test *in vitro* for interactions between PomX variants and PomZ, we used PomZ ATPase activity as a readout. First, we established a base-line for these analyses. The amount of hydrolyzed ATP increased linearly with His_6_-PomZ concentration in the absence of DNA (specific activity: 7±1 ATP h^-1^, 4µM His_6_-PomZ) (Fig. 2e), supporting that His_6_-PomZ hydrolyzes ATP non-cooperatively and is a slow ATPase similar to other ParA-like ATPases. In the presence of increasing concentrations of non-specific herring sperm DNA, His_6_-PomZ ATP hydrolysis was stimulated, reaching saturation at ∼40µg/ml DNA. At this concentration, His_6_-PomZ ATP hydrolysis was stimulated 2-fold (specific activity: ∼15 ATP h^-1^, 4µM His_6_-PomZ) (Fig. 2f). In the presence of saturating concentrations of DNA (60µg/ml), His_6_-PomZ ATPase activity increased linearly with concentration, indicating that PomZ ATPase activity is also non-cooperative in the presence of DNA (Fig. 2e). For comparison, in an average *M. xanthus* cell with one chromosome, the DNA concentration is ∼3400µg/ml suggesting that PomZ *in vivo* works under fully DNA-saturating conditions.

PomX^WT^-His_6_ only stimulated His_6_-PomZ ATPase activity in the presence of DNA (Fig. 2g). Stimulation increased linearly with increasing PomX^WT^-His_6_ concentrations (specific activity: 44±7 ATP h^-1^ at 15µM PomX^WT^-His6, 4µM His_6_-PomZ), and His_6_-PomZ ATPase activity did not reach a plateau even at the highest PomX^WT^-His_6_ concentration (Fig. 2g). Importantly, and in agreement with the BACTH analysis, PomX^N^-His_6_ stimulated His_6_-PomZ ATP hydrolysis in the presence of DNA as efficiently as PomX^WT^-His_6_ (specific activity: 43±4 ATP h^-1^ at 15µM PomX^N^-His_6_, 4µM His_6_-PomZ) (Fig. 2h). By contrast, PomX^C^-His_6_ did not stimulate His_6_-PomZ ATPase activity (Fig. 2i).

Altogether, we conclude that (1) PomX consists of two domains with distinct functions that are both required for PomX activity *in vivo*; (2) PomX^N^ interacts with PomZ and harbors the entire AAP activity; (3) PomX^C^ is required and sufficient to mediate PomX self-interaction with spontaneous filament formation *in vitro* and PomX-dependent cluster incorporation *in vivo*; and, (4) PomX^C^ interacts with PomY.

### Two positively charged residues in PomX^NPEP^ are important for division site and PomX/Y/Z cluster positioning at midcell

To define the PomX^N^ region involved in AAP activity, we performed a detailed sequence analysis of the N-terminal domain of PomX homologs. This analysis revealed a stretch of highly conserved amino acids at the N-terminus (residues 1-22, from hereon PomX^NPEP^) that is enriched in charged amino acids, six of which are positively charged (Fig. 1b, 3a; Supplementary Fig. 3).

**Fig. 3.**
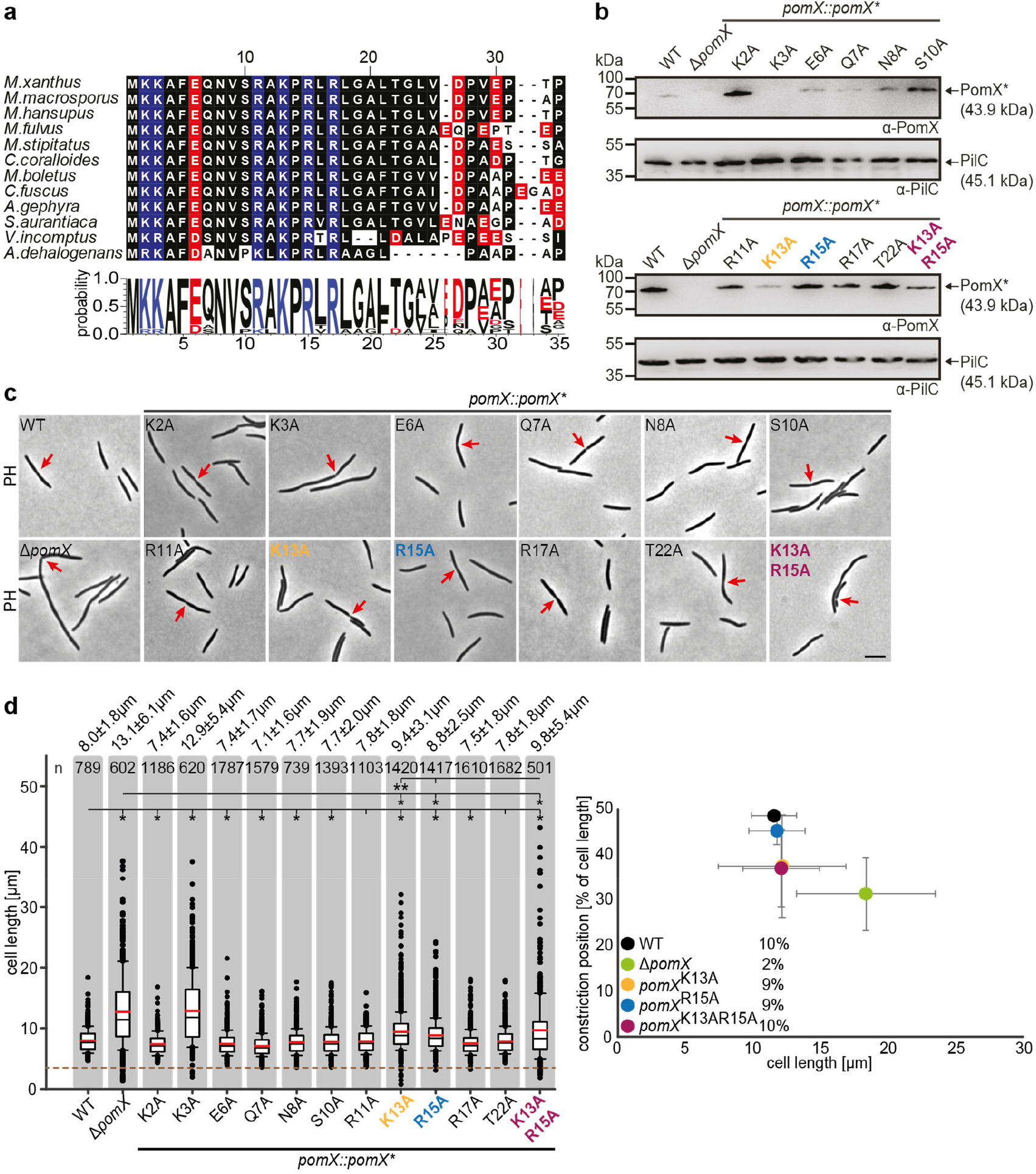
PomX^N^ harbors a conserved N-terminal peptide crucial for cell division site positioning at midcell. **a** Multiple sequence alignment of the conserved PomX N-terminus. Black background indicates similar amino acids. Positively and negatively charged residues are indicated on blue and red, respectively. Weblogo consensus sequence is shown below. **b** Western blot analysis of accumulation of PomX variants. Protein from the same number of cells was loaded per lane. Molecular mass markers are indicated on the left. PilC was used as a loading control. **c** Phase-contrast microscopy of strains of indicated genotypes. Representative cells are shown. Red arrows indicate cell division constrictions. Scale bar, 5µm. **d** Analysis of cell length distribution and cell division constrictions of cells of indicated genotypes. Left panel, boxplot is as in Fig. 1d. * p<0.001; ** p<0.05 in Mann-Whitney test. Right panel, cell division position in % of cell length, and as a function of cell length. Numbers indicate cell division constriction frequency. Number of cells analysed is indicated on top. In b, c and d similar results were obtained in two independent experiments.

To probe the role of PomX^NPEP^, we generated PomX variants lacking residues 2-21 (PomX^Δ2-21^) with and without mCh; however, none of these variants accumulated in *M. xanthus*. Therefore, we performed Ala scanning of PomX^NPEP^ in which all charged or hydrophilic residues were replaced by Ala. Then the mutant alleles replaced the *pomX*^WT^ allele at the *pomX* locus. All 11 PomX variants except for PomX^K3A^ accumulated similarly to or at slightly lower or higher levels than PomX^WT^ (Fig. 3b). Most strains had a cell length similar to WT and cell division constrictions at midcell (Fig. 3c, d). As expected, *pomX*^K3A^ cells were similar to Δ*pomX* cells. More importantly, the *pomX*^K13A^ and *pomX*^R15A^ mutants generated filamentous cells and minicells, but the filamentous cells were shorter than Δ*pomX* cells (Fig. 3d). *pomX*^K13A^ and *pomX*^R15A^ cells had a cell division constriction frequency similar to WT (Fig. 3d), but these constrictions were mostly not at midcell (Fig. 3c, d). The *pomX*^K13AR15A^ double mutant had a more pronounced filamentation phenotype than the single mutants, formed minicells, had a constriction frequency similar to WT, and mostly with the constrictions away from midcell (Fig. 3c, d). PomX^K13AR15A^ accumulated similarly to PomX^WT^ (Fig. 3b). We conclude that Lys13 and Arg15 in PomX^NPEP^ are important for PomX function. Because the PomX^K13A^, PomX^R15A^, and PomX^K13AR15A^ variants caused defects distinct from the Δ*pomX* mutant, i.e., cells are shorter, and with more constrictions, we conclude that substitution of Lys13 and/or Arg15 does not result in a complete loss of PomX function.

mCh-PomX^K13A^, mCh-PomX^R15A^, and mCh-PomX^K13AR15A^ accumulated at the same level as mCh-PomX^WT^ in *M. xanthus* (Supplementary Fig. 4a). All three variants formed clusters *in vivo*; however, these were mostly not at midcell and in the case of mCh-PomX^K13AR15A^ ∼50% localized in the DNA-free subpolar regions while this localization pattern was not observed for mCh-PomX^WT^ (Fig. 4a; Supplementary 4b). The clusters formed by the mutant mCh-PomX variants, similar to those of mCh-PomX^WT^, colocalized with cell division constrictions. Overall, these observations demonstrate that mCh-PomX^K13A^, mCh-PomX^R15A^, and mCh-PomX^K13AR15A^ are functional in forming clusters, defining the division site, and stimulating division but cannot correctly position the division site at midcell. All other PomX variants with substitutions in PomX^NPEP^, including mCh-PomX^K3A^, accumulated and localized as mCh-PomX^WT^ (Supplementary Fig. 4a, c).

**Fig. 4.**
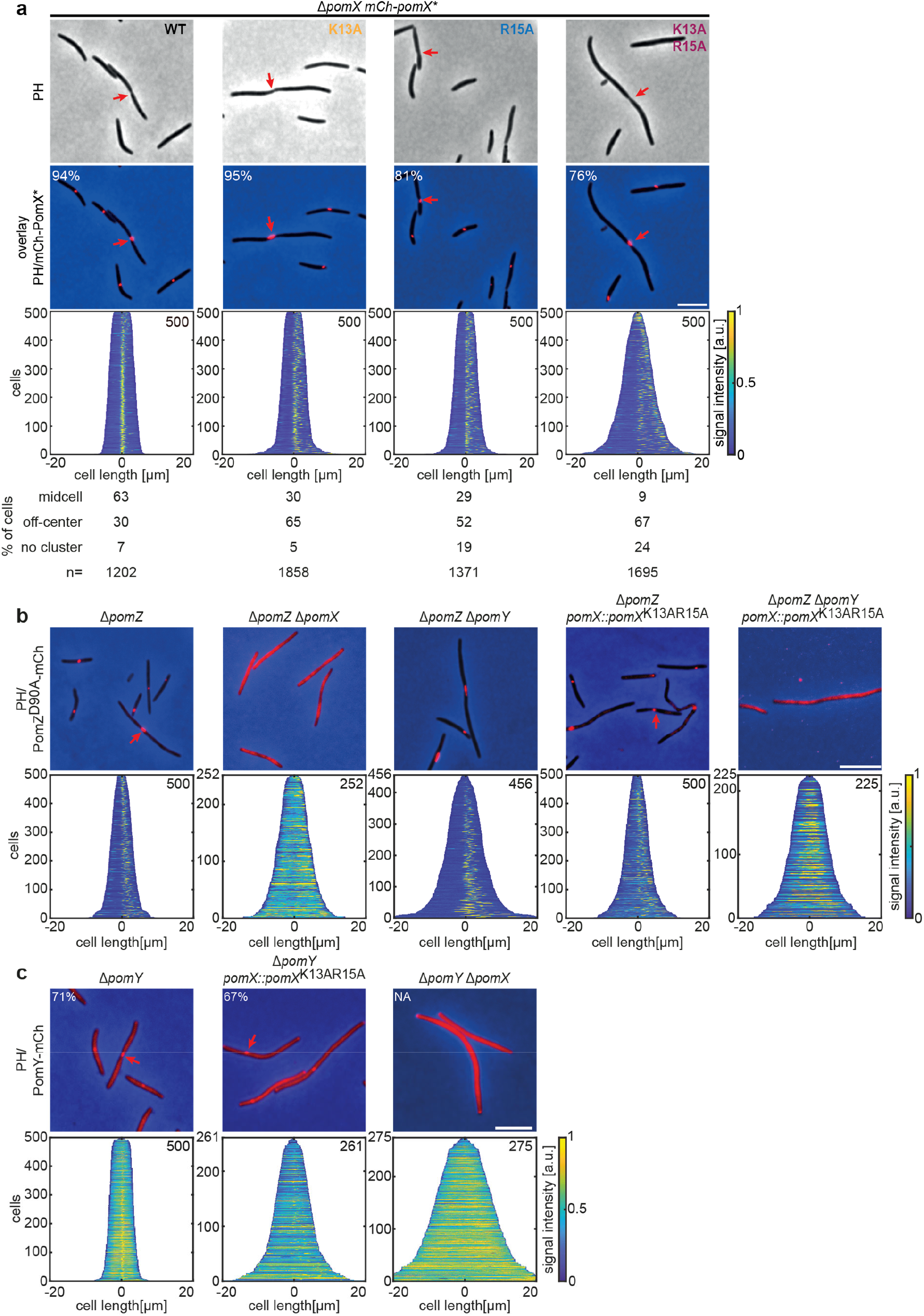
PomX^K13AR15A^ forms clusters and interacts with PomY but not with PomZ *in vivo*. a-c Fluorescence microscopy of cells of indicated genotypes. Phase-contrast (PH) images and/or overlays of fluorescence images and PH of representative cells. Red arrows indicate division constrictions. Scale bar, 5µm. In a, numbers in overlays indicate fraction of cells with a cluster and numbers below indicate localization patterns in % and number of cells analyzed. Demographs are as in Fig. 1e. Similar results were observed in two independent experiments.

Because recruitment of the PomX/Y cluster to the nucleoid and PomX/Y/Z cluster localization at midcell depend on PomZ, these observations pointed in the direction that the PomZ and PomX interaction involves Lys13 and Arg15. To test for an interaction between PomZ and PomX^NPEP^ variants *in vivo*, we explored PomX^K13AR15A^ in more detail and took advantage of the ATP-locked PomZ^D90A^ variant. In the presence of PomX^WT^, PomX/Y/Z^D90A^ clusters are randomly positioned on the nucleoid and rarely at midcell due to lack of PomZ ATPase activity ^32^. We confirmed that in the absence of PomX, PomZ^D90A^-mCh did not form clusters and instead colocalized with the nucleoid, while PomZ^D90A^-mCh still formed clusters in the absence of PomY (Fig. 4b; Supplementary Fig. 5a). In the presence of PomX^K13AR15A^, PomZ^D90A^-mCh formed clusters (Fig. 4b). Because PomZ also interacts with PomY, we speculated that this cluster incorporation resulted from PomY recruiting PomZ^D90A^-mCh. Indeed, upon additional deletion of *pomY*, PomZ^D90A^-mCh no longer formed clusters and colocalized with the nucleoid when PomX^K13AR15A^ was the only source of PomX (Fig. 4b).

An active PomY-mCh fusion did not form clusters in the absence of PomX but formed clusters in the presence of PomX^K13AR15A^ (Fig. 4c; Supplementary Fig. 5b). As expected, in the presence of PomX^K13AR15A^, the PomY-mCh clusters generally localized away from midcell as opposed to clusters in the presence of PomX^WT^.

Altogether, these observations support that PomX^NPEP^ is required for the interaction between PomX and PomZ but for neither PomX self-interaction nor PomX interaction with PomY. Moreover, they support that the PomX/Y/Z clusters formed in the presence of PomX^K13AR15A^ are proficient in stimulating cell division but deficient in efficiently localizing to midcell, consistent with the PomX/PomZ interaction being perturbed. Finally, they support that PomZ in its ATP-bound dimeric form can be recruited independently by PomX and PomY to the PomX/Y complex.

### PomX^NPEP^ is required and sufficient for stimulation of PomZ ATP hydrolysis

We generated PomX constructs for BACTH analysis that were either truncated for PomX^NPEP^ (PomX^Δ2-21^ and PomX^N_Δ2-21^) or contained substitutions in PomX^NPEP^ (PomX^K13AR15A^ and PomX^N_K13AR15A^). PomX^K13AR15A^ and PomX^Δ2-21^ self-interacted, and also interacted with PomX^WT^ and PomY (Fig. 5a). Consistently, *in vitro* PomX^K13AR15A^-His_6_ formed filaments that were bundled by PomY-His_6_, accumulated in the pellet fraction after high-speed centrifugation, and brought PomY-His_6_ to the pellet fraction similarly to PomX^WT^-His_6_ (Fig. 5b, c). As expected, in the BACTH neither PomX^N_Δ2-21^ nor PomX^N_K13AR15A^ interacted with PomX^WT^ and PomY (Fig. 5a). Importantly, all four PomX^NPEP^ mutants were dramatically reduced in interaction with PomZ and PomZ^D90A^ (Fig. 5a). Altogether, these findings further support that PomX^NPEP^ is specifically important for the PomX/PomZ interaction and not for PomX/PomX and PomX/PomY interactions.

**Fig. 5.**
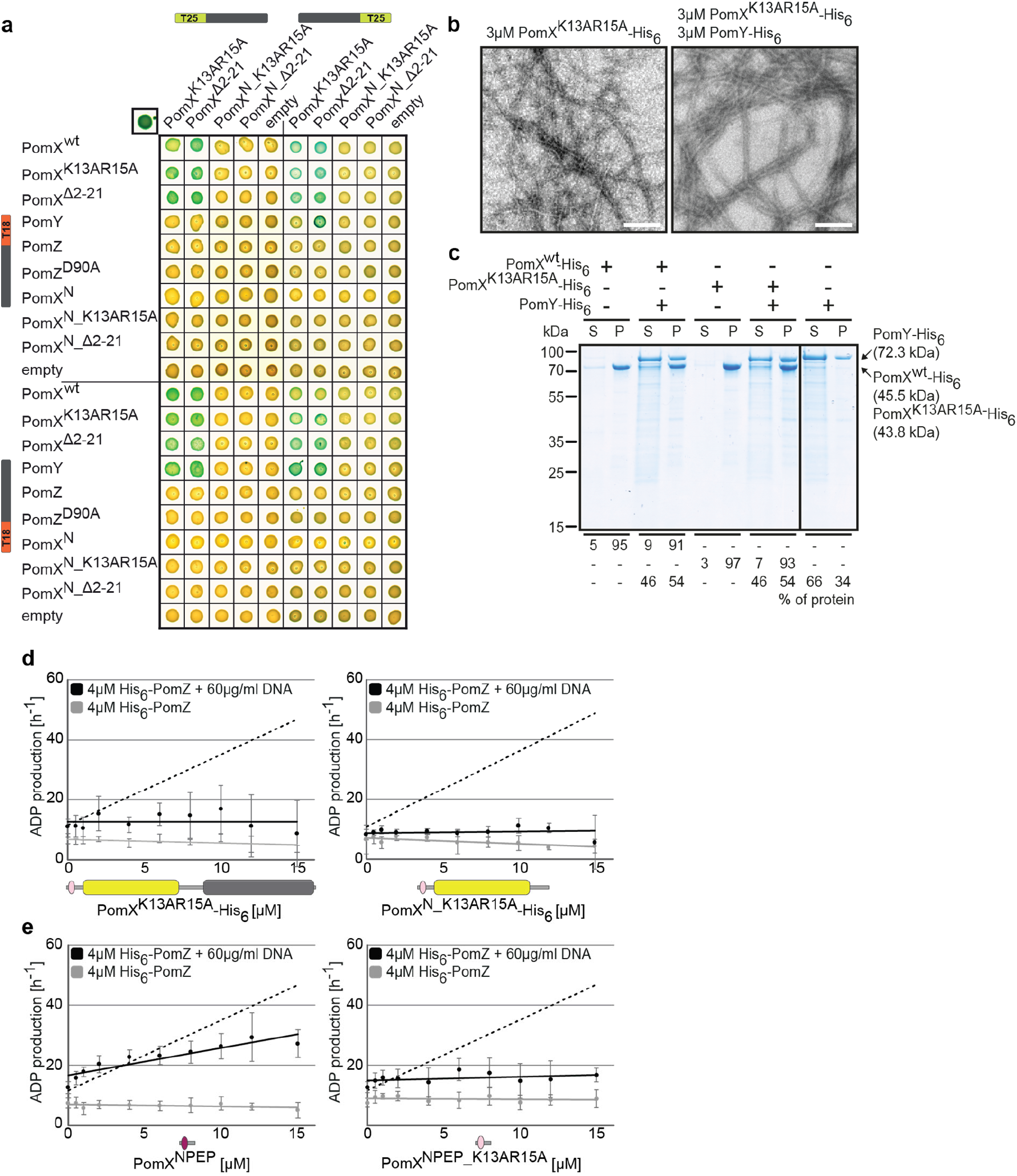
PomX AAP activity resides in PomX^NPEP^. **a** BACTH analysis of interactions between Pom proteins and PomX variants. Experiments were performed as in Fig. 2a. Images show representative results and similar results were obtained in three independent experiments. **b** TEM images of negatively stained purified proteins. Experiments were done as in Fig. 2b. Scale bar, 200nm. Images show representative results of several independent experiments. **c** Sedimentation assays with indicated purified proteins. Proteins were analysed at a concentration of 3µM alone or in combination. After high-speed centrifugation, proteins in the supernatant (S) and pellet (P) fractions were separated by SDS-PAGE and stained with Instant Blue™. Molecular size markers are shown on the left and analysed proteins on the right including their calculated MW. Numbers below indicate % of proteins in different fractions. Similar results were obtained in two independent experiments. All samples were analysed on the same gel; the black line indicates that lanes were removed for presentation purposes. **d, e** His_6_-PomZ ATPase activity. Experiments were done and analysed as in Fig. 2e-i in the presence or absence of DNA and the indicated proteins and peptides. Data points show the mean±STDEV calculated from six independent measurements. In **d**, stippled lines indicates the regression of the ADP production rate in the presence of PomX^WT^-His_6_ (left, Fig. 2g) and PomX^N^-His_6_ (right, Fig 2h). In e, the stippled line indicates the regression of the ADP production rate in the presence of PomX^WT^-His_6_.

Next, we tested whether PomX^NPEP^ is important for PomX AAP activity. Because His_6_-tagged truncated PomX^NPEP^ variants could not be overexpressed in *E. coli*, we focused on the PomX^K13AR15A^-His_6_ and PomX^N_K13RAR15A^-His_6_ variants (Supplementary Fig. 2a). PomX^N_K13AR15A^-His_6_ (calculated MW: 24.1 kDa) behaved similarly to PomX^N^-His_6_ in SEC and eluted as a single peak corresponding to a globular protein of ∼136 kDa (Supplementary Fig. 2b). Remarkably, neither PomX^K13AR15A^-His_6_ nor PomX^N_K13AR15A^-His_6_ stimulated PomZ ATPase activity (Fig. 5d). Consequently, we tested whether a peptide consisting of the 22 PomX^NPEP^ residues alone is sufficient to stimulate His_6_-PomZ ATPase activity. The PomX^NPEP^ peptide alone stimulated His_6_-PomZ ATPase activity in the presence of DNA (specific activity: 27±5 ATP h^-1^ at 15µM PomX^Npep^, 4µM His_6_-PomZ) while a PomX^NPEP^ peptide with the K13AR15A substitutions did not (Fig. 5e). We conclude that PomX^NPEP^ is required and sufficient for stimulation of PomZ ATPase activity by PomX.

### The PomX/PomZ interaction is important for PomX/Y/Z cluster fission during division

24% of cells containing mCh-PomX^K13AR15A^ lacked a visible cluster compared to only 6% in the presence of mCh-PomX^WT^ (Fig. 4a). Tagged and untagged PomX^K13AR15A^ accumulate at the same level as tagged and untagged PomX^WT^ (Fig. 3b: Supplementary Fig. 4a), suggesting that this difference in cluster formation is not caused by differences in gene expression or protein stability. We, therefore, investigated whether mCh-PomX^K13AR15A^ causes a cluster fission defect during cell division.

In the presence of mCh-PomX^WT^, ∼80% of divisions are accompanied by symmetric or asymmetric cluster fission, with each portion of a divided cluster segregating to a daughter (Fig. 6a, b). In the remaining ∼20%, cluster splitting did not visibly occur, the undivided cluster segregated to one of the daughters, and “empty” daughter cells eventually regenerated a cluster that was visible after ∼2h (Fig. 6a). mCh-PomX^K13AR15A^ clusters showed the same three patterns during division. However, cluster fission and segregation to daughters occurred in only ∼20% of cells (Fig. 6a, b).

**Fig. 6.**
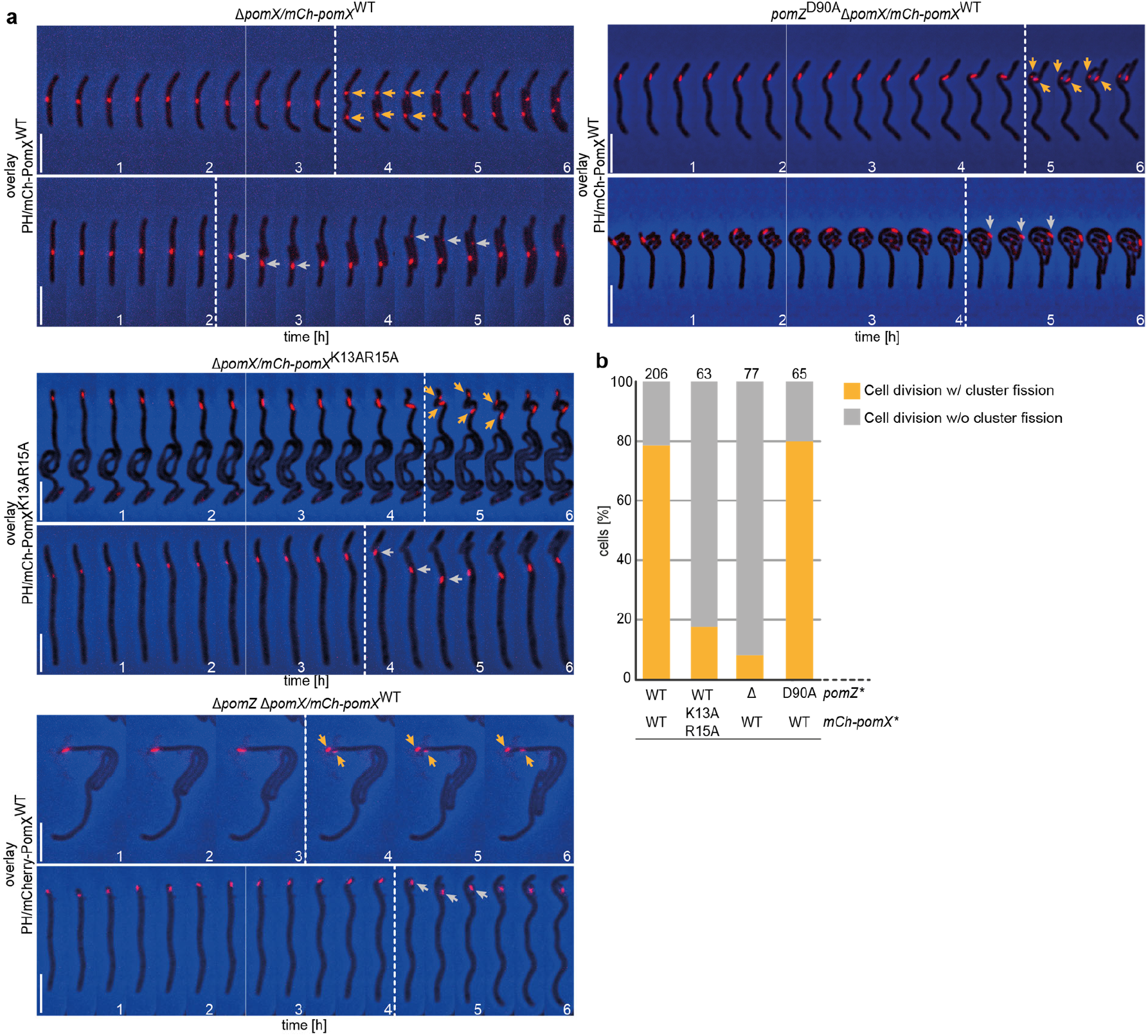
The PomX/PomZ interaction is important for cluster fission during division. **a** Fluorescence time-lapse microscopy of mCh-PomX variants in cells of indicated genotypes. Overlays of representative mCh images and PH are shown in 20 min intervals. Stippled lines indicate cell division events. Orange and grey arrows mark mCh-PomX clusters in daughter cells after cell division with cluster fission and without cluster fission, respectively. Scale bar, 5µm. **b** Quantification of cluster fission during cell division in cells of indicated genotypes. Cell division events were divided into those with (orange) and without (grey) cluster fission. Number of analyzed cell divisions is shown on top. The same results were obtained in two independent experiments.

Because PomX^K13AR15A^ has reduced PomZ AAP activity, we tested cluster fission in Δ*pomZ* cells and in cells containing PomZ^D90A^. In Δ*pomZ* cells, division occurs at a reduced frequency but still over the PomX/Y cluster^32^. In these cells, mCh-PomX^WT^ clusters rarely underwent fission and mostly segregated into one daughter (Fig. 6a, b). In cells with PomZ^D90A^, divisions also occurred over the cluster, and ∼80% of clusters underwent fission during division and segregated to both daughters (Fig. 6a, b). Altogether, these observations suggest that the interaction between PomZ and PomX is important for PomX/Y/Z cluster fission, while ATP hydrolysis by PomZ is not.

## Discussion

In the present study, we used *in vivo* and *in vitro* approaches to functionally dissect the cell division regulatory protein PomX. We demonstrate that PomX is composed of two domains and has three functions *in vivo*. The N-terminal PomX^N^ domain contains the PomZ AAP activity, the C-terminal PomX^C^ domain is essential for PomX self-interaction and PomY interaction and functions as a scaffold for PomX/Y/Z cluster formation, and the PomX/Z interaction, but not PomZ ATPase activity, is important for PomX/Y/Z cluster fission during division.

PomX^N^, which is Ala/Pro-rich and, therefore, likely unstructured, interacts with PomZ and activates ATPase activity of DNA-bound PomZ *in vitro* as efficiently as PomX^WT^. Moreover, a peptide comprising the N-terminal 22 residues of PomX (PomX^NPEP^) is sufficient to activate PomZ ATPase activity. In PomX^NPEP^, two positively charged residues (Lys13 and Arg15) are essential for AAP activity. Of note, the AAP activity of PomX^NPEP^ was lower than that of PomX^N^. Similarly, in the case of the ParB homologs Spo0J of *Thermus thermophilus* and SopB of plasmid F, as well as MinE of *Neisseria gonorrhoeae*, peptides comprising the N-terminal 20, 52, and 22 residues, respectively are sufficient for AAP activity^6,35,36^. In the case of the shorter Spo0J and MinE peptides, the AAP activity was lower than for the full-length proteins^6,36^ while the longer SopB peptide was as efficient as full-length SopB^35^. Because BACTH analyses did not reveal an interaction between PomZ and PomX variants lacking PomX^NPEP^ or containing the K13AR15A substitutions, we speculate that the lower AAP activity of PomX^NPEP^ indicates that the context of PomX^NPEP^ might be important for its AAP activity; however, we cannot rule out that interactions between PomZ and PomX^N^ beyond N^PEP^ may also be important for PomZ ATPase activation.

PomX^C^ with its predicted coiled-coil domain self-interacts and also interacts with PomX^WT^ and PomY. Specifically, PomX^C^, similarly to PomX^WT^, formed filaments *in vitro* that were bundled by PomY. *In vivo* mCh-tagged PomX^C^ integrated into clusters containing PomX^WT^ but alone was not sufficient to form a cluster. By contrast, PomX^N^ did not form or integrate into a cluster under any conditions tested; moreover, PomX^N^ interacted with neither PomX^WT^ nor PomY in BACTH or *in vitro*. Because PomX^WT^ alone can form a cluster and is essential for PomX/Y/Z cluster formation *in vivo*, these observations support a model whereby PomX^WT^ serves as a scaffold protein for cluster formation *in vivo* and in which PomX^C^ has a key role in this scaffolding function by self-interacting and interacting with PomY. The observation that neither PomX^N^ nor PomX^C^ alone is sufficient to form a cluster *in vivo* suggests that these two domains may also interact. Our experiments suggest that these interactions are of low affinity because there were not detected by any of the methods used here (BACTH, TEM, high-speed centrifugation, and pull-down experiments). Alternatively, the conformation of the two separated domains could be different from that in PomX^WT^, and, therefore, no interactions were detected. Altogether, we suggest that PomX^WT^ monomers *in vivo* interact via their coiled-coil domain in PomX^C^ to form a polymeric structure that is stabilized or modified by PomX^N^; this polymeric structure, in turn, recruits PomY. PomZ in its ATP-bound dimeric form is recruited to this structure and associates it to the nucleoid. Because PomX^WT^ and the likely unstructured PomX^N^ stimulate PomZ ATPase activity non-cooperatively and to the same extent, these observations also suggest that PomX^WT^ monomers in the polymeric structure *in vitro* and cluster *in vivo* stimulate PomZ ATPase activity independently of each other.

By analyzing PomZ variants, we previously showed that PomZ ATPase activity is essential for PomX/Y/Z cluster translocation and cluster localization to midcell. Because the PomX variants with reduced AAP activity resulted in the formation of PomX/Y/Z clusters that were typically not at midcell, we conclude that the low intrinsic PomZ ATPase activity is not sufficient to fuel translocation of the PomX/Y/Z cluster to midcell and that this translocation is fueled by PomX, and possibly also by PomY, stimulated ATP hydrolysis by PomZ. In the PomX AAP mutants, constrictions were formed at WT frequencies over the PomX/Y/Z cluster. Because these clusters were typically not at midcell, these PomX variants resulted in divisions away from midcell giving rise to filamentous cells and minicells. Thus, PomX AAP mutants are fully competent in stimulating cell division but deficient in positioning the PomX/Y/Z cluster at midcell. These observations also support that PomX-stimulated PomZ ATPase activity is not important for stimulation of Z-ring formation. Importantly, in mutants lacking PomZ, the PomX/Y cluster is also mostly away from midcell, and Z-rings and constrictions are formed over the cluster away from midcell, but at a much-reduced frequency compared to WT. Thus, PomX AAP mutants and a mutant lacking PomZ phenocopy each other with respect to PomX/Y/Z cluster positioning but not with respect to constriction frequency. We suggest that this difference is the result of different cluster compositions, i.e. the Pom clusters formed in the AAP mutants contain PomZ as well as PomY, which both interact with FtsZ ^32^, while the Pom clusters formed in the absence of PomZ only contain PomX and PomY. Thus, the division defects in Δ*pom* mutants are a consequence of reduced formation and mislocalization of division constrictions, while the PomX AAP mutants are only deficient in cell division localization.

DNA binding by PomZ is important for the low intrinsic as well as PomX-stimulated ATPase activity. This is similar to what has been described for several DNA-binding ParA ATPases and their partner AAPs^6,9,20,32,35,37^. Similarly, MinE-dependent stimulation of ATP hydrolysis by a MinD variant that does not bind the membrane is reduced^24^. However, it remains unknown how DNA or membrane binding makes these ATPases competent for ATPase activity. Likewise, the precise molecular mechanism by which AAPs stimulate ATPase activity of their cognate ParA/MinD family ATPase remains unknown. Nevertheless, our data support that this activation may involve a shared mechanism: In agreement with the findings here that PomX^NPEP^ is enriched in positively charged residues and that Lys13 and Arg15 are essential for PomX AAP activity, it has previously been noted that ParB-type AAPs, MinE-type AAPs and the non-homologous AAP ParG of plasmid TP228 that functions together with the ParA ATPase ParF contain a stretch of N-terminal residues rich in positively charged amino acids ^6,35,36,38^ (Fig. 7). For several of these AAPs, it has been shown that one or more of the positively charged residues are important for AAP activity (Fig. 7)^6,9,21,30,35,36,38^. Also, as noted above, in the case of Spo0J of *T. thermophilus*, plasmid F SopB, and MinE of *N. gonorrhoeae*, 20, 52, and 22 N-terminal residues are sufficient for AAP activity. Thus, PomX is the fourth type of ParA/MinD AAP that displays this characteristic N-terminus and in which positively charged residues are important for AAP activity. Intriguingly, TlpT, which is the suggested AAP of the ParA-like ATPase PpfA involved in cytoplasmic chemoreceptor cluster positioning in *Rhodobacter sphaeroides*^39^, and McdB, which is the AAP of the ParA-like ATPase McdA important for carboxysome positioning in *Synechococcus elongatus*^40^ both have an N-terminal extension enriched in positively charged residues (Fig. 7). Altogether, these findings lend further support to the notion that AAPs of ParA/MinD ATPases use a the same mechanism to stimulate ATPase activity^6,23,38,41^. Moreover, they support that ParA/MinD AAPs display remarkable plasticity and modularity in which a stretch of N-terminal amino acids enriched in positively charged residues grafted onto an interaction domain, which is involved in protein-protein, protein-DNA, or protein-membrane interaction, can generate an AAP.

**Fig. 7.**
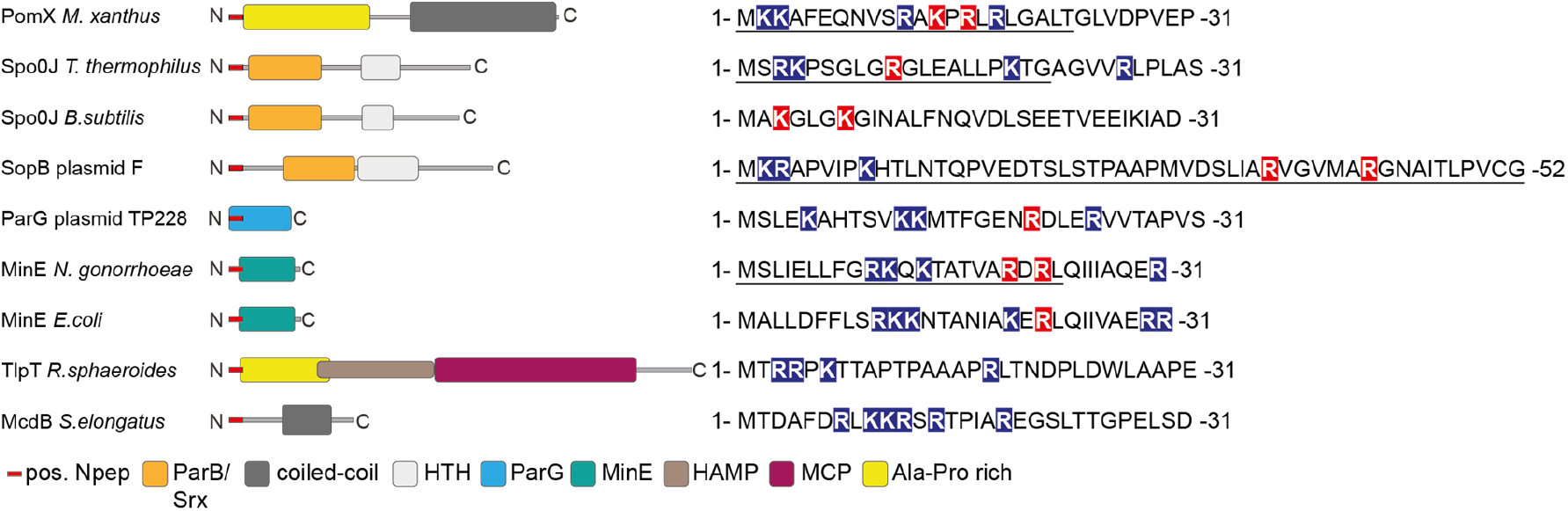
AAPs of MinD/ParA ATPases are diverse but share common features. Left, domain analysis of known and predicted AAPs of ParA/MinD ATPases with key below. Sequences on the right, N-terminus of indicated proteins. Positively charged amino acids are indicated on blue, and positively charged residues experimentally demonstrated to be important for AAP activity on red. Underlined sequences indicate peptides experimentally demonstrated to have AAP activity. Spo0J of *T. thermophilus* and *B. subtilis*, and SopB of plasmid F are ParB homologs.

In addition to serving as a scaffold for PomX/Y/Z cluster formation and as a PomZ AAP, we report that PomX has a third important function in PomX/Y/Z cluster fission during division. Specifically, a PomX AAP mutant did not support fission as efficiently as PomX^WT^. Similarly, in cells lacking PomZ, the PomX/Y cluster did not split efficiently during division. By contrast, ATP-locked dimeric PomZ^D90A^ stimulated fission as efficiently as PomZ^WT^. In all these mutants, division occurs over the Pom cluster. Additionally, in cells treated with cephalexin, which blocks cell division, cluster fission does not occur ^32,33^. Because cephalexin treated cells still segregate their chromosomes, these observations support that cell division is essential for fission, chromosome segregation is not sufficient, and the PomX/Z interaction is important while PomZ ATPase is not. In the future, it will be interesting to establish the mechanism for PomX/Y/Z cluster fission in detail.

## Methods

### *M. xanthus* and *E. coli* strains and growth

Strains, plasmids and primers are listed in Supplementary Table 1, 2 and 3, respectively. *M. xanthus* strains are derivatives of DK1622 WT^42^. *M. xanthus* strains were cultivated in 1% CTT medium (1% casitone, 10 mM Tris-HCl pH 7.6, 1 mM KPO_4_ pH 7.6, 8 mM MgSO_4_) or on 1% CTT 1.5% agar plates^43^. Kanamycin, oxytetracycline, and gentamycin were added at concentrations of 50 µg/ml, 10 µg/ml, and 10 µg/ml, respectively. Growth was measured as an increase in optical density (OD) at 550nm. *M. xanthus* cells were transformed by electroporation. In-frame deletions were generated as described^44^. Plasmids were integrated by site-specific recombination at the Mx8 *attB* locus or by homologous recombination at the endogenous site. All plasmids were verified by sequencing. All strains were verified by PCR. Non-motile strains were generated to allow time-lapse microscopy for several hours by deletion of *mglA*^45,46^. *E. coli* strains were grown in LB or 2xYT medium in the presence of relevant antibiotics or on LB plates containing 1.5% agar^47^. Plasmids were propagated in *E. coli* NEB Turbo cells (New England Biolabs) (F’ *proA*_^*+*^_*B*^*+*^ *lacI*^*q*^ *ΔlacZM15/fhuA2 Δ(lac-proAB) glnV galK16 galE15 R(zgb-210::Tn10)* Tet^S^ *endA1 thi-1 Δ(hsdS-mcrB)5*). Growth of *E. coli* was measured as an increase in OD at 600nm.

### Fluorescence microscopy and live cell imaging

Fluorescence microscopy was performed as described^32^. Briefly, exponentially growing cells were transferred to slides with a thin 1.0% agarose pad (SeaKem LE agarose, Cambrex) with TPM buffer (10 mM Tris-HCl pH 7.6, 1 mM KH_2_PO_4_ pH 7.6, 8 mM MgSO_4_), covered with a coverslip and imaged using a temperature-controlled Leica DMi6000B inverted microscope with an HCX PL FLUOTAR objective at 32°C. Phase-contrast and fluorescence images were recorded with a Hamamatsu ORCA-flash 4.0 sCMOS camera using the LASX software (Leica Microsystems). Time-lapse microscopy was performed as described^46^. Briefly, cells were transferred to a coverslip mounted on a metallic microscopy slide and covered with a pre-warmed 1% agarose pad supplemented with 0.2% casitone in TPM buffer. Slides were covered with parafilm to retain humidity of the agarose. Live-cell imaging was performed at 32°C. For DNA staining, cells were incubated with 1 mg/ml 2-(4-Amidinophenyl)-6-indolcarbamidine-dihydrochloride (DAPI) for 10 min at 32°C before microscopy. Image processing was performed with Metamorph_v 7.5 (Molecular Devices). For image analysis, cellular outlines were obtained from phase-contrast images using Oufti and manually corrected if necessary^48^. Fluorescence microscopy image analysis was performed with a custom-made Matlab script (Matlab R2018a, MathWorks). To identify clusters of mCh-PomX in cells, all pixels above a threshold intensity, given by four standard deviations above the mean cytosolic fluorescence, were determined. If more than 20 pixels with intensities above the threshold were connected (assuming connectivity of 4), they were identified as a cluster. Demographs were generated by randomly selecting 500 cells from a dataset (unless otherwise stated) and calculating the length of the cells’ centerlines (using meshes obtained from Oufti). Next, the fluorescence intensity of each cell was corrected by the background fluorescence locally around each cell. Cells were ordered by length and oriented according to the cell segment with the brightest intensity. The brightest 3% of the intensity values of all cells were set to the maximal intensity value, which is scaled to one, to be able to visualize fluorescence intensity variations inside the cells despite very bright clusters.

### Bacterial Two Hybrid Assay

BACTH experiments were performed as described^49^. Relevant genes were cloned into the appropriate vectors to construct N-terminal and C-terminal fusions with the 25-kDa N-terminal or the 18-kDa C-terminal adenylate cyclase fragments of *B. pertussis*. Restoration of cAMP production was observed by the formation of blue color on LB agar supplemented with 80 µg/ml 5-bromo-4-chloro-3-indolyl-β-D-galactopyranoside (X-Gal) and 0.25 mM isopropyl-β-D-thiogalactoside (IPTG). As positive control, the leucine zipper from GCN4 was fused to the T18 and the T25 fragment. As a negative control, plasmids were co-transformed that only expressed the T18 or T25 fragment. All tested interactions were spotted on the same LB agar plates with positive control and all corresponding negative controls.

### Western blot analysis

Western blot analyses were performed as described^47^ with rabbit polyclonal α-PomX (1:10000), α-PomY (1:10000), α-PomZ (1:5000)^32^, α-PilC (1:3000)^50^ or α-mCh (1:5000; Biovision) primary antibodies together with horseradish-conjugated goat α-rabbit immunoglobulin G (Sigma-Aldrich) as secondary antibody. Blots were developed using Luminata Forte Western HRP Substrate (Millipore) and visualized using a LAS-4000 luminescent image analyzer (Fujifilm).

### Protein purification

His_6_-PomZ was purified from *E. coli* as described^33^. PomX^WT^-His_6_ and PomY-His_6_ were purified as described^32^ using plasmids pEMR3 and pEMR1, respectively. PomX^K13AR15A^-His_6_ was purified as PomX-His_6_. Briefly, plasmid pSH58 was propagated in *E. coli* NiCo21(DE3) cells (NEB), grown in LB medium with 50µg/ml kanamycin at 30°C to an OD_600_ of 0.6-0.7. Protein expression was induced with 0.4mM IPTG for 16h at 18°C. Cells were harvested by centrifugation at 6000*g* for 20min at 4°C. Cells were washed with lysis buffer 1 (50mM NaH_2_PO_4_; 300mM NaCl; 10mM imidazole; pH 8.0 (adjusted with NaOH)) and lysed in 50ml lysis buffer 2 (lysis buffer 1 with 0.1mM EDTA; 1mM β-mercaptoethanol; 100mg/ml phenylmethylsulfonyl fluoride (PMSF); 1× complete protease inhibitor (Roche Diagnostics GmbH); 10U/ml DNase 1) by sonication in 3 rounds of sonication for 5min with a Branson Sonifier (Duty cycle 4; output control 40%) (Heinemann) on ice. Cell debris was removed by centrifugation at 4,700*g* for 45min at 4°C. PomX^K13AR15A^-His_6_ was affinity purified with Protino Ni-NTA resin (Macherey-Nagel) from a batch, equilibrated in lysis buffer 1. PomX^K13AR15A^-His_6_ was eluted from the resin by washing 1× with 5ml elution buffer 1 (lysis buffer 1 with 50mM imidazole) and 3 x with 5ml elution buffer 2 (lysis buffer 1 with 250mM imidazole). Purified PomX^K13AR15A^-His_6_ was dialyzed 4× against 2l dialysis buffer (50mM Hepes/NaOH pH 7.2; 50mM KCl; 0.1mM EDTA; 1mM β-mercaptoethanol; 10% (v/v) glycerol). Proteins were frozen in liquid nitrogen and stored at -80°C until used.

To purify PomX^N^-His_6_ plasmid pAH157 was propagated in *E. coli* NiCo21(DE3) cells, grown in 2×YT medium with 50µg/ml kanamycin and 0.5% glucose at 30°C to an OD_600_ of 0.6-0.7. Protein expression was induced with 0.4mM IPTG for 16h at 18°C. Cells were harvested and lysed as described for PomX^K13AR15A^. PomX^N^-His_6_ was affinity purified with Protino Ni-NTA resin (Macherey-Nagel) from a batch, equilibrated in lysis buffer 1. Contaminating proteins were eluted from the resin by washing 6× with 40ml wash buffer 1 (lysis buffer 1 with 20mM imidazole), 1× with 40ml wash buffer 2 (lysis buffer 1 with 50mM imidazole). PomX^N^-His_6_ was eluted from the resin by washing with 1× 10 ml elution buffer 1 (lysis buffer 1 with 100mM imidazole), 1× with 10 ml elution buffer 2 (lysis buffer 1 with 150mM imidazole) and 1× with 10ml elution buffer 3 (lysis buffer 1 with 200mM imidazole). The elution fractions were pooled and loaded onto a HiLoad 16/600 Superdex 200 pg gel filtration column (GE Healthcare) that was equilibrated with dialysis buffer. Elution fractions were pooled. Proteins were frozen in liquid nitrogen and stored at -80°C until used.

For purification of PomX^N_K13AR15A^-His_6_ plasmid pAH165 was propagated in *E. coli* Rosetta2(DE3) cells, grown in 2×YT medium with 50µg/ml kanamycin, 30µg/ml chloramphenicol and 0.5% glucose at 30°C to an OD_600_ of 0.6-0.7. Protein expression was induced with 0.5mM IPTG for 16h at 18°C. Cells were harvested and lysed as described for PomX^K13AR15A^. PomX^N_K13AR15A^-His_6_ was purified from cleared lysates with a 5ml HiTrap Chelating HP column loaded with NiSO_4_ and equilibrated with lysis buffer 1. The column was washed with 20 column volumes (CVs) lysis buffer 1. The protein was eluted with elution buffer (50mM NaH_2_PO_4_; 300mM NaCl; 500mM imidazole; pH 8.0 (adjusted with NaOH)) with a gradient of 20 CV. Fractions containing PomX^N_K13AR15A^-His_6_ were pooled and concentrated with an Amicon Ultra-15 centrifugation filter device with a cutoff of 3kDa and loaded onto a HiLoad 16/600 Superdex 200 pg gel filtration column (GE Healthcare) that was equilibrated with dialysis buffer. Elution fractions were pooled. Proteins were frozen in liquid nitrogen and stored at -80°C until used.

To purify PomX^C^-His_6_ plasmid pAH152 was propagated in *E*.*coli* Rosetta2(DE3) cells, grown in 2×YT medium with 50µg/ml kanamycin, 30µg/ml chloramphenicol and 0.5% glucose at 37°C to an OD_600_ of 0.6-0.7. Protein expression was induced with 0.5mM IPTG for 4h at 37°C. Cells were harvested and lysed as described for PomX^K13AR15A^. PomX^C^-His_6_ was affinity purified with Protino Ni-NTA resin (Macherey-Nagel) from a batch, equilibrated with lysis buffer 1. Contaminating proteins were eluted from the resin by washing 6× with 40ml wash buffer 1 (lysis buffer 1 with 20mM imidazole), 1× with 40ml wash buffer 2 (lysis buffer 1 with 50mM imidazole) and 2× with 40ml wash buffer 3 (lysis buffer 1 with 100mM imidazole) and 1× with 40ml wash buffer 4 (lysis buffer 1 with 150mM imidazole). PomX^C^-His_6_ was eluted from the resin with 3× 10ml elution buffer (lysis buffer 1 with 250mM imidazole). The elution fractions were pooled and dialyzed against 4× 2l of dialysis buffer at 4°C. Proteins were frozen in liquid nitrogen and stored at -80°C until used.

For purification of PomX^N^-Strep, plasmid pDS232 was propagated in *E. coli* Rosetta2(DE3) cells, grown in LB medium with 50µg/ml kanamycin, 30µg/ml chloramphenicol and 0.5% glucose at 32°C to an OD_600_ of 0.6-0.7. Protein expression was induced with 0.5mM IPTG for 2h at 32°C. Cells were harvested and lysed as described for PomX^K13AR15A^-His_6_ but in StrepTag lysis buffer (100mM Tris-HCl pH 8.0, 150mM NaCl, 1mM EDTA, 1mM dithiothreitol (DTT)). PomX^N^-Strep was purified from cleared lysates with a 5ml StrepTrap HP column equilibrated with StrepTag lysis buffer. The column was washed with 20 CV StrepTag lysis buffer. The protein was eluted with StrepTag elution buffer (StrepTag lysis buffer with 2.5mM D-desthiobiotin). Fractions containing PomX^N^-Strep were pooled and dialyzed against 2 x 3l of dialysis buffer at 4°C. Proteins were frozen in liquid nitrogen and stored at -80°C until used.

For purification of PomX^C^-Strep, plasmid pDS333 was propagated in *E. coli* Rosetta2(DE3) cells, grown in LB medium with 50µg/ml kanamycin, 30µg/ml chloramphenicol and 0.5% glucose at 32°C to an OD_600_ of 0.6-0.7. Protein expression was induced with 1mM IPTG for 18h at 18°C. Cells were harvested and lysed as described for PomX^N^-Strep. PomX^C^-Strep was purified from a batch using 2ml Strep-TactinXT 4Flow resin (iba), equilibrated with StrepTag lysis buffer. The resin was incubated with the cleared lysate for 1h at 4°C on a rotary shaker. Contaminating proteins were eluted from the resin by washing 5× with 10ml StrepTag lysis buffer. The protein was eluted with 1× BXT buffer (100mM Tris-HCl pH8.0, 150mM NaCl, 1mM EDTA, 50mM biotin) (iba). Fractions containing PomX^C^-Strep were pooled and dialyzed against 2× 5l of dialysis buffer at 4°C. Proteins were frozen in liquid nitrogen and stored at -80°C until used.

### Protein sedimentation assay

Before sedimentation experiments, a clearing spin was performed for all proteins to be analyzed at 20,000 *g* for 10min at 4°C. Proteins at a final concentration of 3µM in a total volume of 50µl were mixed and incubated for 1h at 32°C in a buffer (50mM Hepes/NaOH, pH 7.2, 50mM KCl, 1mM β-mercaptoethanol, 10mM MgCl_2_). Samples were separated into soluble and insoluble fractions by high-speed centrifugation (160,000 *g*, 60min, 25°C). Insoluble and soluble fractions were separated, and volumes adjusted with 1× SDS sample buffer. Fractions were separated by SDS-PAGE and stained with Instant Blue™ (expedion) for 10 min.

### *In vitro* pull-down experiments

10µM protein alone or pre-mixed as indicated were incubated for 1h at 32°C in reaction buffer (50mM HEPES/NaOH pH 7.2, 50mM KCl, 10mM MgCl_2_) in a total volume of 200µl and applied to 20µl 5% (v/v) MagStrepXT beads (iba) for 30min. Magnetic beads were washed 10× with 200µl reaction buffer. Proteins were eluted with 200µl 1× BXT buffer (100mM Tris-HCl pH8.0, 150mM NaCl, 1mM EDTA, 50mM biotin) (iba). 10 µl per sample were separated by SDS-PAGE.

### Negative stain transmission electron microscopy

To fix and stain protein samples for negative stain TEM, 10µl of a protein sample of interest (protein concentration before application onto the EM grid 3µM) was applied onto an EM grid (Plano) and incubated for 1min at 25°C. Residual liquid was blotted through the grid by applying the grid’s unused side on Whatman paper. The grid was washed twice with double-distilled H_2_O. For staining, 10µl of 1% uranyl acetate solution was applied onto the grid for 1min and blotted through with a Whatman paper. If protein mixtures were applied to the EM grid, proteins of interest at a concentration of 3µM were pre-mixed in a low-binding microtube (Sarstedt) and incubated for 10min at 25°C before application onto the EM grid. Finished grids were stored in a grid holder for several months at room temperature. Electron microscopy was performed with a CM120 electron microscope (FEI) at 120kV.

### ATPase assay

ATP hydrolysis was determined using a 96-well NADH-coupled enzymatic assay^51^ with modifications. Protein concentration was determined using Protein Assay Dye Reagent Concentrate (BioRad). Assays were performed in reaction buffer (50mM HEPES/NaOH pH 7.2, 50mM KCl, 10mM MgCl_2_) with 0.5mM nicotinamide adenine dinucleotide (NADH) and 2mM phosphoenolpyruvate and 3µl of a pyruvate kinase/lactate dehydrogenase mix (PYK/LDH; Sigma). PomX^NPEP^ and PomX^NPEP_K13AR15A^ peptides (MKKAFEQNVSRA**K**P**R**LRLGALT and MKKAFEQNVSRA**A**P**A**LRLGALT) were purchased from Thermo Scientific. If appropriate, herring sperm DNA was added at a concentration of 60µg/ml unless otherwise stated. Buffer was pre-mixed with proteins in low-binding microtubes (Sarstedt) on ice. To correct for glycerol in the assays, dialysis buffer was added if necessary. 100µl mixtures were transferred into transparent UV-STAR µCLEAR 96-well microplates (Greiner bio-one). The reaction was started by the addition of 1mM ATP. Measurements were performed in an infinite M200PRO (Tecan) for 2h in 30sec intervals at 32°C shaking at 340nm wavelength. To account for background by spontaneous ATP hydrolysis and UV-induced NADH decomposition, all assays were performed without the addition of His_6_-PomZ and measurements were subtracted. The light path was determined experimentally with known NADH concentrations to be 0.248cm. The extinction coefficient of NADH ε340 = 6220 M^-1^cm^-1^ was used.

### Analytical size-exclusion chromatography

Experiments were carried out in dialysis buffer (50mM Hepes/NaOH pH 7.2; 50mM KCl; 0.1mM EDTA; 1mM β-mercaptoethanol; 10% (v/v) glycerol). PomX^N^-His_6_ and PomX^N_K13AR15A^-His_6_ were applied onto a Superdex 200 10/300GL gel filtration column equilibrated with dialysis buffer. Blue dextran (2000 kDa), ferritin (440 kDa), conalbumin (75 kDa), ovalbumin (43 kDa), carbonic anhydrase (29 kDa), RNAse A (13.7 kDa), and aprotinin (6.5 kDa) were used as standards with the same buffer conditions to calibrate the column.

### Bioinformatics

Gene and protein sequences of PomX, PomY, and PomZ were obtained from NCBI. PomX homologs were identified in a best-best hit reciprocal BlastP analysis from fully-sequenced genomes of Myxobacteria^52^. The similarity and identity of proteins were calculated from pairwise sequence alignments with EMBOSS Needle^53^. Domain analyses were performed with SMART^54^, PROSITE and Pfam^55^. Multiple sequence alignments were created with MUSCLE^53^ and further edited with Bioedit (https://bioedit.software.informer.com/7.2/). Consensus sequences of multiple sequence protein alignments were created with Weblogo 3^56^. Proteins used: PomXMx (ABF89666; MXAN_0636), PomXMm (ATB45064; MYMAC_000648), PomXMh (AKQ69458; A176_006370), PomXMf (AKF79435; MFUL124B02_03625), PomXMs (AGC41991; MYSTI_00641), PomXCc (AFE03552; COCOR_00544), PomXMb (ATB30647; MEBOL_004108), PomXCf (ATB35470; CYFUS_000883), PomXAg (AKI99966; AA314_01593), PomXSa (ADO75429; STAUR_7674), PomXVi (AKU92216; AKJ08_2603), PomXAd (ABC83600; Adeh_3834). Spo0J Tt (AAS81946.1), Spo0J Bs (P26497.2) ParG TP228 Ec (WP_139578510.1), SopB F Ec (BAA97917.1), MinE Ng (AAK30127.1), MinE Ec (EFB7450413.1), TlpT Rs (ABA79218.1), McdB Se (ABB57864.1).

### Statistics

The mean and standard deviation (STDEV) were calculated with Excel 2016. Localization patterns from fluorescence microscopy data were quantified based on the indicated n-value per strain. Boxplots were generated with SigmaPlot 14.0 (Systat). Statistical analysis was performed with SigmaPlot 14.0. All data sets were tested for normality using a Shapiro-Wilk test. For data with a non-normal distribution, a Mann-Whitney test was applied to test for significant differences.

### Data availability

The authors declare that all data supporting this study are available within the article or its Supplementary Information file. The source data underlying Fig. 1a, b, d, 2a, b, c, d, e, f, g, h, i, 3a, b, d, 4a, b, c, 5a, b, c, d, e, 6b, 7 and Supplementary Fig. 1b, 2a, b, c, d, e, f, 4a, b, 5a, b are provided as a Source Data file.

### Code availability

Custom MATLAB scripts used for fluorescence microscopy data analysis is available from the corresponding author upon request.

## Acknowledgements

We thank Sabrina Huneke-Vogt for assistance with plasmid constructions, Manon Wigbers for help with the image analysis script, and Anke Treuner-Lange for many helpful discussions.

## Authors’ contributions

DS: Designed and conceived the study, performed most of the experiments and analysed data.

AH and DS: Purified proteins, performed enzymate assays and protein-protein interaction analyses.

SB: Generated Matlab scripts for analysis of fluorescence microscopy images EF: Supervised research and provided funding.

LSA: Designed and conceived the study, supervised research and provided funding. DS, AH, SB and LSA: Analyzed and interpreted data.

DS and LSA: Wrote the manuscript. All authors approved the final manuscript.

