## Supplementary Information for "PomX, a ParA/MinD ATPase activating protein, is a triple regulator of cell division in *Myxococcus xanthus*"

**This file contains**

- Supplementary Figure S1-S5
- Supplementary Table 1-3
- Supplementary References

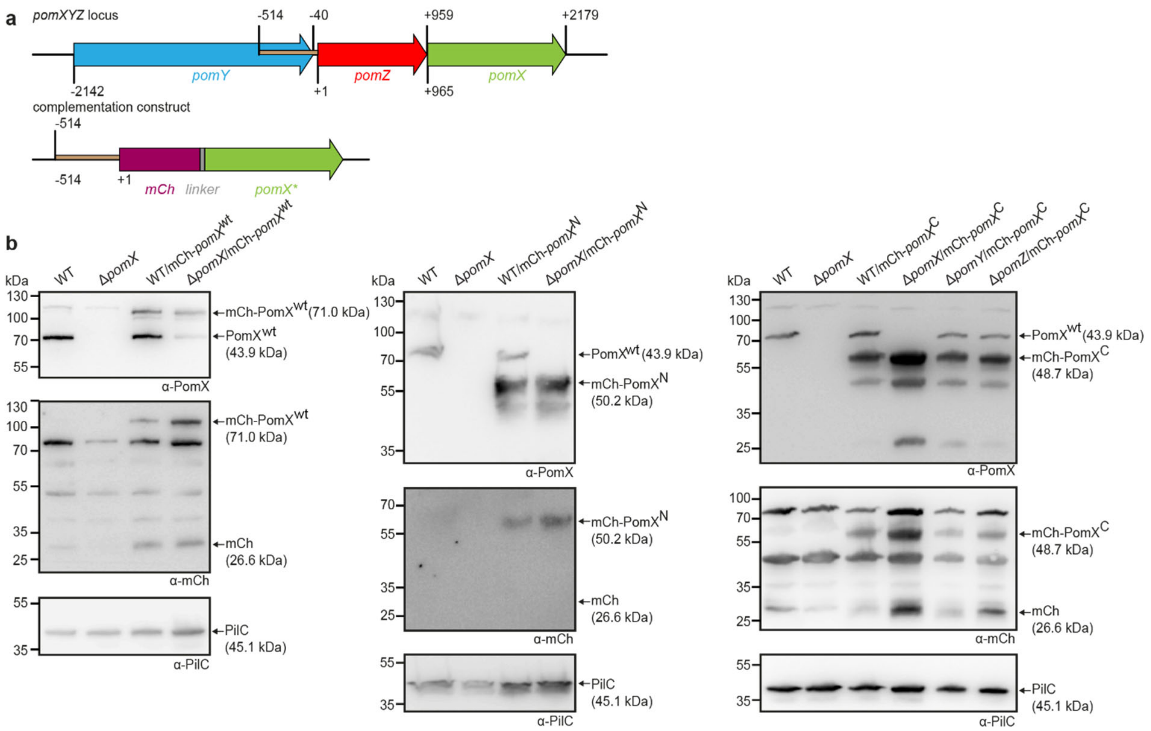


**Supplementrary Fig. 1. PomX variants accumulate in *M.xanthus*.**

**a** Schematic of *pomXYZ* locus (upper panel) and the construct used for ectopic expression of *mCh-pomX* and its variants from the *attB* site (lower panel). The brown region upstream of *pomZ* was used as a promoter for the expression of *mCh-pomX* variants. All coordinates are relative to to first nucleotide in *pomZ* start codon (+1). **b** Western blot analysis of mCh-PomX (71.0 kDa), mCh-PomX$^{N}$ (50.2 kDa), and mCh-PomX$^{C}$ (48.7 kDa) accumulation in indicated strains. Protein from the same number of cells was loaded per lane. Molecular mass markers are indicated on the left and analysed proteins on the right including calculated MW. The same blots were sequentially analyzed with α-PomX (top panel), α-mCh (middle panel), and α-PilC (lower panel). PilC was used as a loading control. Note PomX$^{WT}$ (43.9 kDa) does not migrate at the expected size in SDS-PAGE but as a protein of a molecular weight of ~72 kDa. Similarly, mCh-PomX$^{WT}$, mCh-PomX$^{N}$, and mCh-PomX$^{C}$ migrate at ~110 kDa, ~60 kDa, and ~62 kDa, respectively.

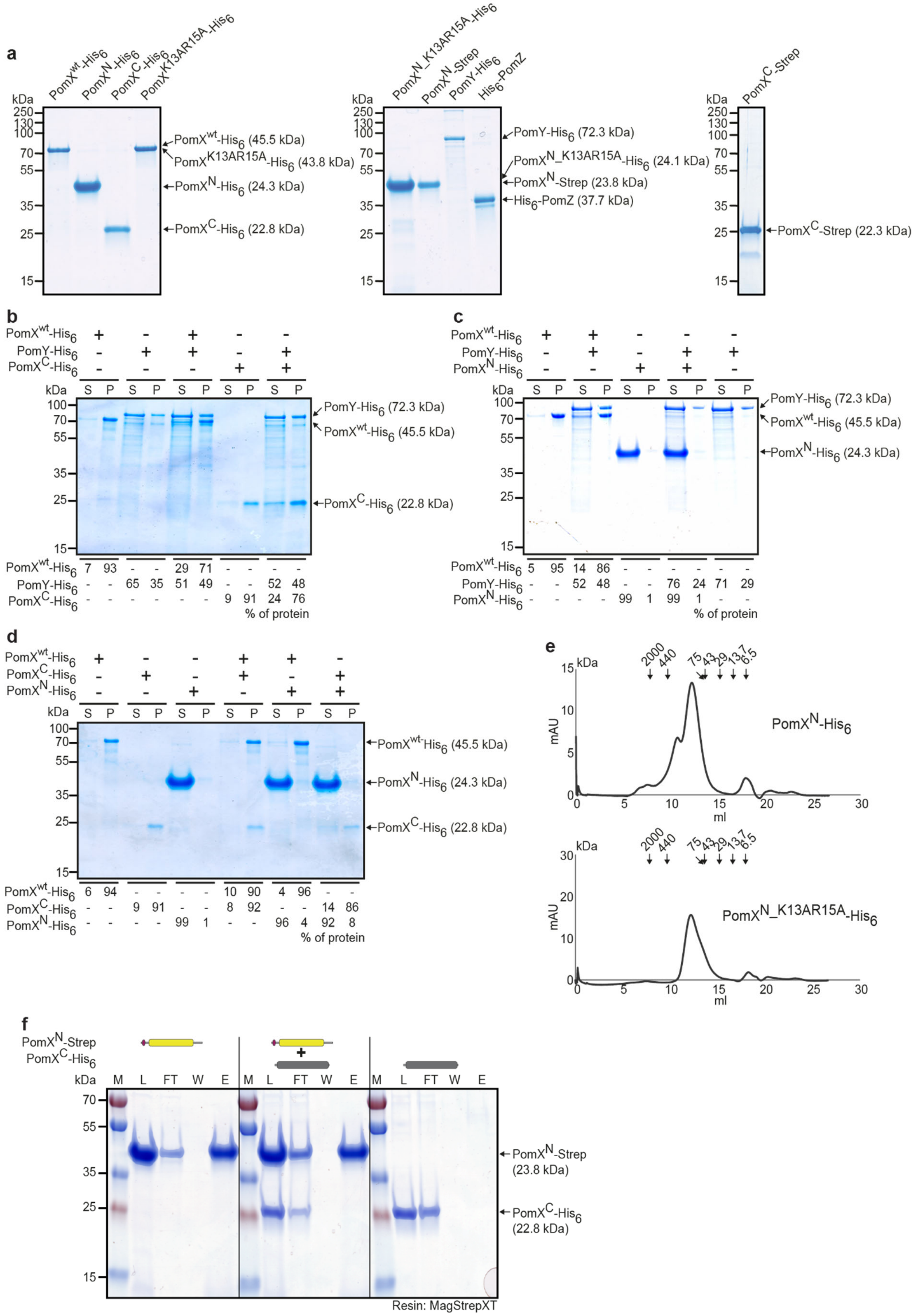


**Supplementrary Fig. 2. Purification and analysis of Pom proteins.**

**a** SDS-PAGE analysis of purified proteins used in this study. Molecular size markers are shown on the left and the purified proteins inclusing calculated MW on the right. 2µg per protein was loaded. Note that $PomX^{wt}$-$His_6$, $PomX^{N}$-$His_6$, $PomX^{K12AR15A}$-$His_6$, $PomX^{N\_K13AR15A}$-$His_6$, and $PomX^{N}$-Strep do not separate according to their calculated MW. **b-d** Sedimentation assays with indicated proteins. The indicated proteins were mixed at final concentrations of 3µM as indicated. Following high-speed ultracentrifugation, the supernatant (S) and pellet )P) fractions were separated by SDS-PAGE. Molecular size markers are shown on the left and analysed proteins on the right. Numbers below show the quantification of indicated protein in the different fractions in %. Similar results were observed in two independent experiments. **e** Size exclusion chromatography elution profile of $PomX^{N}$-$His_6$ and $PomX^{N\_K13AR15A}$-$His_6$. The elution pattern of $PomX^{N}$-$His_6$ and $PomX^{N\_K13AR15A}$-$His_6$ from a Superdex 200 10/300GL gel filtration column was measured at 280nm. Arrows indicate elution maxima of protein standards of the indicated size in kDa. The same results were observed in two independent experiments. **f** *In vitro* pull-down experiments with purified $PomX^{N}$-Strep and $PomX^{C}$-$His_6$. Instant $Blue^{TM}$-stained SDS-PAGE shows load (L), flow-through (FL), wash (W), and elution (E) fractions using MagStrep XT beads in pull-down experiments with 10µM of indicated proteins alone or pre-mixed as indicated on top. Molecular size markers are shown on the left and proteins analysed on the right together with their calculated MW. All samples in a panel were analysed on the same gel and black lines are included for clarity. Experiments were repeated in two independent experiments with similar results.

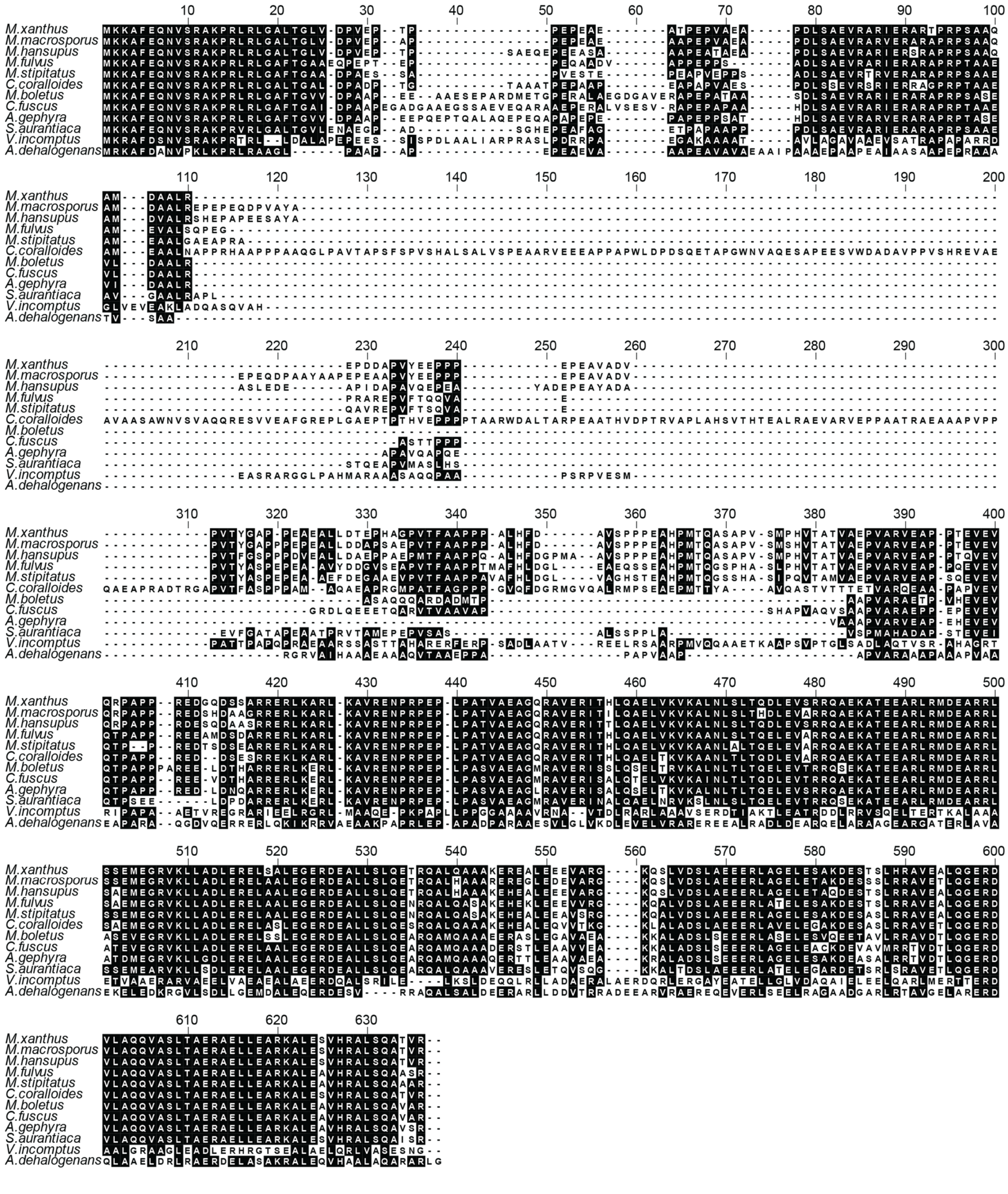


**Supplementrary Fig. 3. PomX homologs are highly conserved.**
Alignment of PomX homologs from other fully sequenced genomes of Myxobacteria. Sequences were aligned with MUSCLE and color-coded by homology using Bioedit. Black and white backgrounds indicate similar/homologous amino acids and no conservation, respectively.

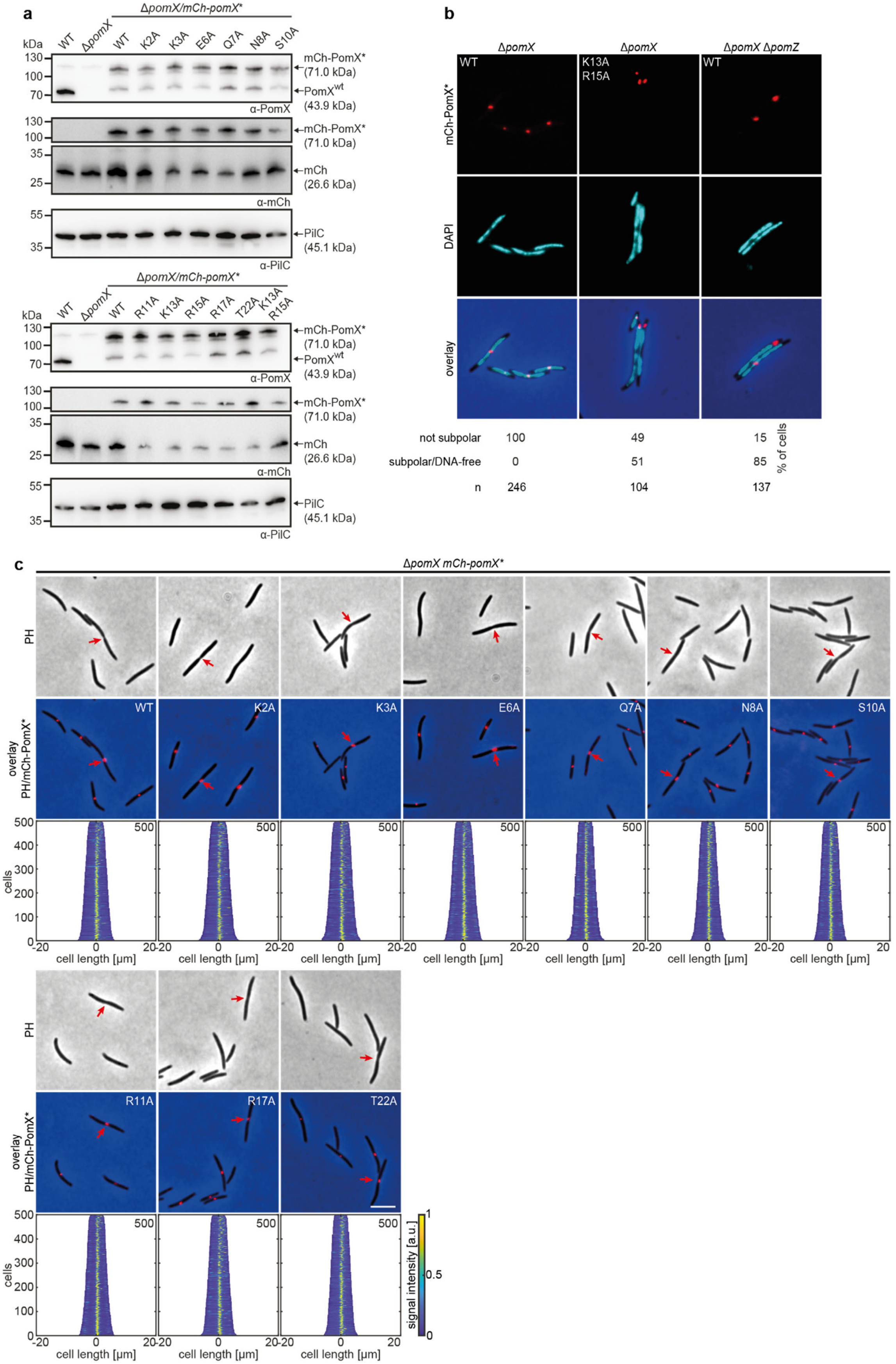


**Supplementrary Fig. 4. The PomX$^{K13AR15A}$ variant is impaired in function.**

**a** Western blot analysis of the accumulation of mCh-PomX variants in indicated strains. Protein from the same number of cells was loaded per lane. Molecular mass marker is shown on the left and analysed proteins on the right. The same blots were sequentially analyzed with α-PomX (top panel), α-mCh (middle panel), and α-PilC (lower panel) antibodies. PilC was used as a loading control. Note PomX (43.9 kDa) does not migrate at the expected size in SDS-PAGE but instead as a protein of a molecular weight of 72 kDa. Similarly, mCh-PomX migrates at ~110 kDa. Similar results were obtained in two independent experiments. **b** Fluorescence microscopy of mCh-PomX variants in DAPI stained cells of the indicated genotype. The mCh signal (first panel), DAPI signal (second panel), and the overlay (third panel) show representative cells. Amino acid substitutions are indicated in white in the mCh images. Scale bar, 5μm. Quantification of mCh-PomX* localization patterns in % and the number of analyzed cells is shown below the images. Images show representative cells. Similar results were obtained in two independent experiments. **c** Fluorescence microscopy of indicated mCh-PomX variants. Phase-contrast and fluorescence images of representative cells and the overlay are shown. Red arrows indicate cell division constrictions. Scale bar, 5μm. Demographs were created as in Fig. 1e. Experiments were repeated in two independent experiments with similar results.

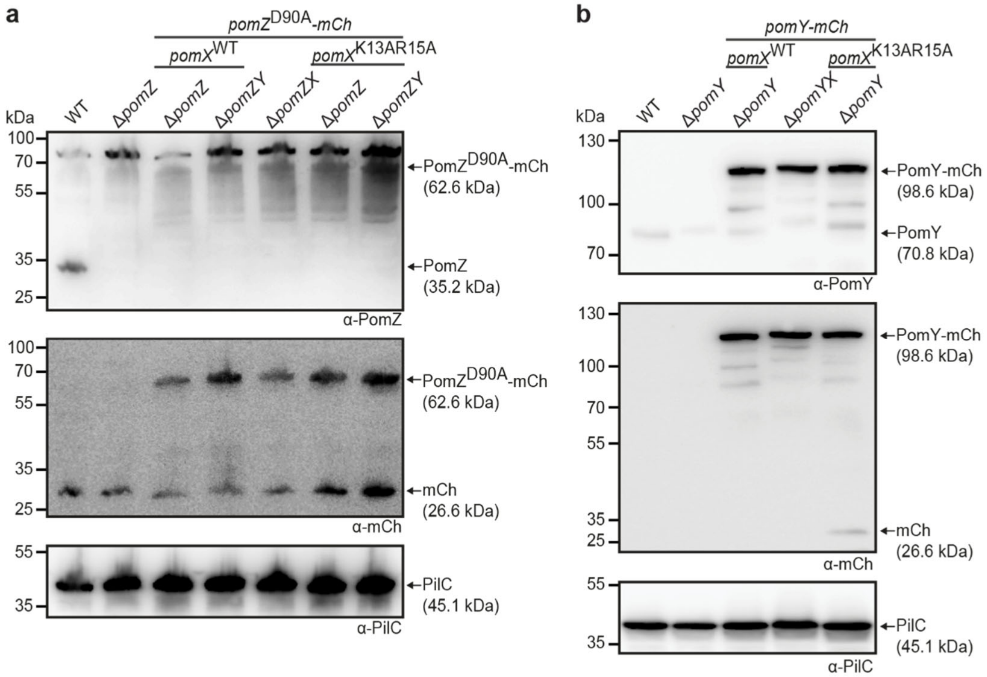


**Supplementary Fig. 5. Western blow analysis of PomY-mCh and PomZ$^{D90A}$-mCh accumulation.**

**a** Western blot analysis of PomZ$^{D90A}$-mCh accumulation in indicated strains. Protein from the same number of cells was loaded per lane. Molecular mass markers are indicated on the left and analysed proteins including MW on the right. The same blots were sequentially analyzed with α-PomZ (top panel), α-mCh (middle panel), and α-PilC (lower panel). PilC was used as a loading control. The same results were observed in two independent experiments. **b** Western blot analysis of PomY-mCh accumulation in indicated strains. Blots were done as in **a**, but α-PomY were used instead of α-PomZ. The same results were observed in two independent experiments.

**Supplementary Table 1.** *M. xanthus* strains used in this study

| **Strain** | **Genotype**[1] | **Source/reference** |
|---|---|---|
| DK1622 | Wild-type | [1] |
| SA3108 | Δ*pomZ* | [2] |
| SA3146 | Δ*pomZ*; *attB*::$P_{mxan0635}$ *pomZ*$^{D90A}$*-mCh* (pKA43) | [2] |
| SA4223 | Δ*pomX* | [3] |
| SA4252 | Δ*pomX*; *attB*::$P_{mxan0635}$ *mCh-pomX* (pAH53) | [3] |
| SA4297 | Wild-type; *attB*::$P_{mxan0635}$ *mCh-pomX* (pAH53) | This study |
| SA4703 | Δ*pomY* | [2] |
| SA4712 | Δ*pomY*; *attB*::$P_{pilA}$ *pomY-mCh* (pDS7) | [2] |
| SA4797 | Δ*mglA*; Δ*pomX*; *attB*::$P_{mxan0635}$ *mCh-pomX* (pAH53) | [3] |
| SA6100 | *pomX::pomX*$^{K13AR15A}$ | This study |
| SA7014 | Δ*pomX*; Δ*pomZ*; *attB*::$P_{mxan0635}$ *pomZ*$^{D90A}$*-mCh* (pKA43) | This study |
| SA7061 | Δ*mglA*; Δ*pomZ*; Δ*pomX*; *attB*::$P_{mxan0635}$ *mCh-pomX* (pAH53) | This study |
| SA7063 | Δ*pomZ*; Δ*pomX*; *attB*::$P_{mxan0635}$ *mCh-pomX* (pAH53) | This study |
| SA8240 | *pomX::pomX*$^{K13AR15A}$; Δ*pomZ*; *attB*::$P_{mxan0635}$ *pomZ*$^{D90A}$*-mCh* (pKA43) | This study |
| SA8250 | *pomX::pomX*$^{K13AR15A}$; Δ*pomY*; *attB*::$P_{pilA}$ *pomY-mCh* (pDS7) | This study |
| SA8268 | *pomX::pomX*$^{K13AR15A}$; Δ*pomY*; Δ*pomZ*; *attB*::$P_{mxan0635}$ *pomZ*$^{D90A}$*-mCh* (pKA43) | This study |
| SA9700 | *pomX::pomX*$^{E6A}$ | This study |
| SA9701 | *pomX::pomX*$^{Q7A}$ | This study |
| SA9702 | *pomX::pomX*$^{N8A}$ | This study |
| SA9714 | *pomX::pomX*$^{R11A}$ | This study |
| SA9715 | *pomX::pomX*$^{K3A}$ | This study |
| SA9716 | *pomX::pomX*$^{R17A}$ | This study |
| SA9717 | *pomX::pomX*$^{T22A}$ | This study |
| SA9718 | *pomX::pomX*$^{K2A}$ | This study |
| SA9719 | *pomX::pomX*$^{R15A}$ | This study |
| SA9720 | Δ*pomY*; Δ*pomZ*; *attB*::$P_{mxan0635}$ *pomZ*$^{D90A}$*-mCh* (pKA43) | This study |
| SA9721 | Δ*pomX*; Δ*pomY*; *attB*::$P_{pilA}$ *pomY-mCh* (pDS7) | This study |
| SA9726 | Δ*pomX*; *attB*::$P_{mxan0635}$ *mCh-pomX*$^{N}$ (pDS252) | This study |
| SA9727 | Wild-type; *attB*::$P_{mxan0635}$ *mCh-pomX*$^{N}$ (pDS252) | This study |
| SA9731 | *pomX::pomX*$^{K13A}$ | This study |
| SA9732 | *pomX::pomX*$^{S10A}$ | This study |
| SA9739 | Δ*pomX*; *attB*::$P_{mxan0635}$ *mCh-pomX*$^{Q7A}$ (pDS317) | This study |
| SA9740 | Δ*pomX*; *attB*::$P_{mxan0635}$ *mCh-pomX*$^{N8A}$ (pDS318) | This study |
| SA9741 | Δ*pomX*; *attB*::$P_{mxan0635}$ *mCh-pomX*$^{R17A}$ (pDS323) | This study |
| SA9742 | Δ*pomX*; *attB*::$P_{mxan0635}$ *mCh-pomX*$^{T22A}$ (pDS324) | This study |
| SA9743 | Δ*pomX*; *attB*::$P_{mxan0635}$ *mCh-pomX*$^{S10A}$ (pDS319) | This study |
| SA9744 | Δ*pomX*; *attB*::$P_{mxan0635}$ *mCh-pomX*$^{R11A}$ (pDS320) | This study |
| SA9747 | Δ*pomX*; *attB*::$P_{mxan0635}$ *mCh-pomX*$^{K13A}$ (pDS321) | This study |
| SA9748 | Δ*pomX*; *attB*::$P_{mxan0635}$ *mCh-pomX*$^{R15A}$ (pDS322) | This study |
| SA9749 | Δ*pomX*; *attB*::$P_{mxan0635}$ *mCh-pomX*$^{K2A}$ (pDS314) | This study |
| SA9750 | Δ*pomX*; *attB*::$P_{mxan0635}$ *mCh-pomX*$^{K3A}$ (pDS315) | This study |
| SA9751 | Δ*pomX*; *attB*::$P_{mxan0635}$ *mCh-pomX*$^{E6A}$ (pDS316) | This study |
| SA9752 | Δ*pomX*; *attB*::$P_{mxan0635}$ *mCh-pomX*$^{K13AR15A}$ (pDS325) | This study |
| SA9753 | Δ*mglA*; Δ*pomX*; *attB*::$P_{mxan0635}$ *mCh-pomX*$^{K13AR15A}$ (pDS325) | This study |
| SA9754 | Δ*mglA*; Δ*pomZ*; Δ*pomX*; $P_{mxan0635}$ *mCh-pomX* (pAH53); *mxan18-19::*$P_{mxan0635}$ *pomZ*$^{D90A}$ (pDS80) | This study |

| | | |
|---|---|---|
| SA9755 | Wild-type; *attB*::$P_{mxan0635}$ *mCh-pomX*$^{C}$ (pDS329) | This study |
| SA9756 | Δ*pomY*; *attB*::$P_{mxan0635}$ *mCh-pomX*$^{C}$ (pDS329) | This study |
| SA9757 | Δ*pomZ*; *attB*::$P_{mxan0635}$ *mCh-pomX*$^{C}$ (pDS329) | This study |
| SA9762 | Δ*pomX*; *attB*::$P_{mxan0635}$ *mCh-pomX*$^{C}$ (pDS329) | This study |

[1] Plasmids in brackets encode indicated genes or gene fusions and are integrated in a single copy at indicated sites in the genome. Genes expressed from the *attB* site or the *mxan18-19* intergenic region were expressed from the *pilA* promoter ($P_{pilA}$) or the native promoter of *pomZ* ($P_{mxan0635}$).

**Supplementary Table 2.** Plasmids used in this study

| Plasmid | Description[1] | Source/reference |
|---|---|---|
| pAH27 | Construct for in-frame deletion of *pomX,* KmR | [3] |
| pAH53 | $P_{mxan0635}$ *mCh-pomX,* Mx8 *attB,* KmR | [3] |
| pAH152 | Overexpression of $PomX^{C}$-$His_6$, KmR | This study |
| pAH154 | $P_{mxan0635}$ *mCh-*$pomX^{N}$*,* Mx8 *attB,* KmR | This study |
| pAH157 | Overexpression of $PomX^{N}$-$His_6$, KmR | This study |
| pAH165 | Overexpression of $PomX^{N\_K13AR15A}$-$His_6$, KmR | This study |
| pDS1 | Construct for in-frame deletion of *pomY,* KmR | [3] |
| pDS7 | $P_{pilA}$ *pomY-mCh,* Mx8 *attB,* KmR | This study |
| pDS16 | Construct for in-frame deletion of *pomY* & *pomZ,* KmR | This study |
| pDS80 | $P_{mxan0635}$ $pomZ^{D90A}$, *mxan_18-19* intergenic region*,* TcR | This study |
| pDS100 | BACTH plasmid for *pomZ* (pUT18C), AmpR | This study |
| pDS103 | BACTH plasmid for *pomX* (pUT18C), AmpR | This study |
| pDS106 | BACTH plasmid for *pomX* (pKT25), KmR | This study |
| pDS109 | BACTH plasmid for *pomZ* (pUT18), AmpR | This study |
| pDS110 | BACTH plasmid for *pomX* (pUT18), AmpR | This study |
| pDS114 | BACTH plasmid for *pomX* (pKNT25) KmR | This study |
| pDS115 | BACTH plasmid for $pomZ^{D90A}$ (pUT18C), AmpR | This study |
| pDS117 | BACTH plasmid for $pomZ^{D90A}$ (pUT18), AmpR | This study |
| pDS120 | BACTH plasmid for *pomY* (pUT18C), AmpR | This study |
| pDS122 | BACTH plasmid for *pomY* (pUT18), AmpR | This study |
| pDS184 | BACTH plasmid for $pomX^{\Delta 2\text{-}21}$ (pUT18), AmpR | This study |
| pDS185 | BACTH plasmid for $pomX^{\Delta 2\text{-}21}$ (pUT18C), AmpR | This study |
| pDS186 | BACTH plasmid for $pomX^{\Delta 2\text{-}21}$ (pKT25), KmR | This study |
| pDS187 | BACTH plasmid for $pomX^{\Delta 2\text{-}21}$ (pKNT25) KmR | This study |
| pDS188 | BACTH plasmid for $pomX^{C}$ (pUT18), AmpR | This study |
| pDS189 | BACTH plasmid for $pomX^{C}$ (pUT18C), AmpR | This study |
| pDS190 | BACTH plasmid for $pomX^{C}$ (pKT25), KmR | This study |
| pDS191 | BACTH plasmid for $pomX^{C}$ (pKNT25) KmR | This study |
| pDS192 | BACTH plasmid for $pomX^{N}$ (pUT18), AmpR | This study |
| pDS193 | BACTH plasmid for $pomX^{N}$ (pUT18C), AmpR | This study |
| pDS194 | BACTH plasmid for $pomX^{N}$ (pKT25), KmR | This study |
| pDS195 | BACTH plasmid for $pomX^{N}$ (pKNT25) KmR | This study |
| pDS232 | Overexpression of $PomX^{N}$-Strep, KmR | This study |
| pDS252 | $P_{mxan0635}$ *mCh-*$pomX^{N}$*,* Mx8 *attB,* KmR | This study |
| pDS253 | BACTH plasmid for $pomX^{N\_K13AR15A}$ (pUT18), AmpR | This study |
| pDS254 | BACTH plasmid for $pomX^{N\_K13AR15A}$ (pUT18C), AmpR | This study |
| pDS255 | BACTH plasmid for $pomX^{N\_K13AR15A}$ (pKT25), KmR | This study |
| pDS256 | BACTH plasmid for $pomX^{N\_K13AR15A}$ (pKNT25) KmR | This study |
| pDS257 | BACTH plasmid for $pomX^{N\_\Delta 2\text{-}21}$ (pUT18), AmpR | This study |
| pDS258 | BACTH plasmid for $pomX^{N\_\Delta 2\text{-}21}$ (pUT18C), AmpR | This study |
| pDS259 | BACTH plasmid for $pomX^{N\_\Delta 2\text{-}21}$ (pKT25), KmR | This study |
| pDS260 | BACTH plasmid for $pomX^{N\_\Delta 2\text{-}21}$ (pKNT25) KmR | This study |
| pDS303 | nat. site codon exchange for $pomX^{K2A}$, KmR | This study |
| pDS304 | nat. site codon exchange for $pomX^{K3A}$, KmR | This study |
| pDS305 | nat. site codon exchange for $pomX^{E6A}$, KmR | This study |
| pDS306 | nat. site codon exchange for $pomX^{Q7A}$, KmR | This study |

| | | |
|---|---|---|
| pDS307 | nat. site codon exchange for *pomX*$^{N8A}$, KmR | This study |
| pDS308 | nat. site codon exchange for *pomX*$^{S10A}$, KmR | This study |
| pDS309 | nat. site codon exchange for *pomX*$^{R11A}$, KmR | This study |
| pDS310 | nat. site codon exchange for *pomX*$^{K13A}$, KmR | This study |
| pDS311 | nat. site codon exchange for *pomX*$^{R15A}$, KmR | This study |
| pDS312 | nat. site codon exchange for *pomX*$^{R17A}$, KmR | This study |
| pDS313 | nat. site codon exchange for *pomX*$^{T22A}$, KmR | This study |
| pDS314 | $P_{mxan0635}$ *mCh-pomX*$^{K2A}$, Mx8 *attB,* KmR | This study |
| pDS315 | $P_{mxan0635}$ *mCh-pomX*$^{K3A}$, Mx8 *attB,* KmR | This study |
| pDS316 | $P_{mxan0635}$ *mCh-pomX*$^{E6A}$, Mx8 *attB,* KmR | This study |
| pDS317 | $P_{mxan0635}$ *mCh-pomX*$^{Q7A}$, Mx8 *attB,* KmR | This study |
| pDS318 | $P_{mxan0635}$ *mCh-pomX*$^{N8A}$, Mx8 *attB,* KmR | This study |
| pDS319 | $P_{mxan0635}$ *mCh-pomX*$^{S10A}$, Mx8 *attB,* KmR | This study |
| pDS320 | $P_{mxan0635}$ *mCh-pomX*$^{R11A}$, Mx8 *attB,* KmR | This study |
| pDS321 | $P_{mxan0635}$ *mCh-pomX*$^{K13A}$, Mx8 *attB,* KmR | This study |
| pDS322 | $P_{mxan0635}$ *mCh-pomX*$^{R15A}$, Mx8 *attB,* KmR | This study |
| pDS323 | $P_{mxan0635}$ *mCh-pomX*$^{R17A}$, Mx8 *attB,* KmR | This study |
| pDS324 | $P_{mxan0635}$ *mCh-pomX*$^{T22A}$, Mx8 *attB,* KmR | This study |
| pDS325 | $P_{mxan0635}$ *mCh-pomX*$^{K13AR15A}$, Mx8 *attB,* KmR | This study |
| pDS329 | $P_{mxan0635}$ *mCh-pomX*$^{C}$, Mx8 *attB,* KmR | This study |
| pDS333 | Overexpression of PomX$^{C}$-Strep, KmR | This study |
| pEMR1 | Overexpression of PomY-$His_6$, KmR | This study |
| pEMR3 | Overexpression of PomX-$His_6$, KmR | [3] |
| pKA1 | Construct for in-frame deletion of *pomZ,* KmR | [2] |
| pKA3 | Overexpression of $His_6$-PomZ, KmR | [2] |
| pKA43 | $P_{mxan0635}$ *pomZ*$^{D90A}$*-mCh*, Mx8 *attB,* TcR | [2] |
| pMAT12 | Construct for in-frame deletion of *pomZ* & *pomX,* KmR | [3] |
| pSH1 | nat. site codon exchange for *pomX*$^{K13AR15A}$, KmR | This study |
| pSH36 | BACTH plasmid for *pomX*$^{K13AR15A}$ (pKNT25) KmR | This study |
| pSH37 | BACTH plasmid for *pomX*$^{K13AR15A}$ (pKT25), KmR | This study |
| pSH38 | BACTH plasmid for *pomX*$^{K13AR15A}$ (pUT18), AmpR | This study |
| pSH39 | BACTH plasmid for *pomX*$^{K13AR15A}$ (pUT18C), AmpR | This study |
| pSH58 | Overexpression of PomX$^{K13AR15A}$-$His_6$, KmR | This study |
| pSL16 | Construct for in-frame deletion of *mglA,* KmR | [4] |
| pUT18 | BACTH plasmid | [5] |
| pUT18C | BACTH plasmid | [5] |
| pKT25 | BACTH plasmid | [5] |
| pKNT25 | BACTH plasmid | [5] |

[1] Genes expressed from the *attB* site or the *mxan18-19* intergenic region were expressed from the *pilA* promoter ($P_{pilA}$) or the native promoter of *pomZ* ($P_{mxan0635}$).

**Supplementary Table 3.** Primers used in this study

| **Primer** | **Sequence 5'-3'** [1] |
|---|---|
| pomX BTH fwd XbaI | GCGTCTAGAGATGAAGAAAGCCTTTGAAC |
| pomX BTH rev KpnI | GCGGGTACCCGGCGCACCGTGGCCTGAC |
| pomY BTH fwd XbaI | GCGTCTAGAGGTGAGCGACGAGCGTCCG |
| pomY BTH rev KpnI | GCGGGTACCCGAGCGGCGAAGTATTTGTG |
| pomZ BTH fwd XbaI | GCGTCTAGAGATGGAAGCGCCGACGTAC |
| pomZ BTH rev KpnI | GCGGGTACCCGGCCGGCCTGCTGGGTGCC |
| pomXΔ2-21 BTH fwd XbaI | GCGTCTAGAGATGACGGGCCTCGTCGACCCC |
| pomXC BTH fwd XbaI | GCGTCTAGAGATGGCCACCGTGGCGGAGGCG |
| pomXN BTH rev KpnI | GCGGGTACCCGGGGCAGCGGCTCCGGGCG |
| 0636 up fwd | GCGGGATCCGTCACCCCAAGCCATTC |
| PomX K2A rev native | CAAAGGCTTTCGCCATGGTTCTCAG |
| PomX K2A fwd native | CTGAGAACCATGGCGAAAGCCTTTG |
| 0636 HindIII rev stop | GCGAAGCTTTCAGCGCACCGTGGCCTGAC |
| PomX K3A rev native | CTGTTCAAAGGCCGCCTTCATGGTTC |
| PomX K3A fwd native | GAACCATGAAGGCGGCCTTTGAACAG |
| PomX E6A rev | GGACACGTTCTGCGCAAAGGCTTTCTT |
| PomX E6A fwd | AAGAAAGCCTTTGCGCAGAACGTGTCC |
| PomX Q7A rev | GCGGGACACGTTCGCTTCAAAGGCTTT |
| PomX Q7A fwd | AAAGCCTTTGAAGCGAACGTGTCCCGC |
| PomX N8A rev | GGCGCGGGACACCGCCTGTTCAAAGGC |
| PomX N8A fwd | GCCTTTGAACAGGCGGTGTCCCGCGCC |
| PomX S10A rev | CGGCTTGGCGCGCGCCACGTTCTGTTC |
| PomX S10A fwd | GAACAGAACGTGGCGCGCGCCAAGCCG |
| PomX R11A rev | GCGCGGCTTGGCCGCGGACACGTTCTG |
| PomX R11A fwd | CAGAACGTGTCCGCGGCCAAGCCGCGC |
| PomX K13A rev | GCGGAGGCGCGGCGCGGCGCGGGACAC |
| PomX K13A fwd | GTGTCCCGCGCCGCGCCGCGCCTCCGC |
| PomX R15A rev | GCCCAGGCGGAGCGCCGGCTTGGCGCG |
| PomX R15A fwd | CGCGCCAAGCCGGCGCTCCGCCTGGGC |

| | |
|---|---|
| PomX R17A rev | CAGCGCGCCCAGCGCGAGGCGCGGCTT |
| PomX R17A fwd | AAGCCGCGCCTCGCGCTGGGCGCGCTG |
| PomX T22A rev | GTCGACGAGGCCCGCCAGCGCGCCCAG |
| PomX T22A fwd | CTGGGCGCGCTGGCGGGCCTCGTCGAC |
| PomX K13AR15A rev | CAGAACGTGTCCCGCGCCGCGCCGGCCCTCCGCCTGGGCGCGCTG |
| PomX K13AR15A fwd | CAGCGCGCCCAGGCGGAGGGCCGGCGCGGCGCGGGACACCTTCTG |
| mCherry XbaI fwd | GCGTCTAGAGTGAGCAAGGGCGAGGAG |
| PomX K2A rev | TTCAAAGGCTTTCGCCATGGCTCCGCC |
| PomX K2A fwd | GGCGGAGCCATGGCGAAAGCCTTTGAA |
| KA348 | GCCAAGCTTTCAGCGCACCGTGGCCTG |
| PomX K3A fwd | GGAGCCATGAAGGCGGCCTTTGAACAG |
| PomX K3A rev | CTGTTCAAAGGCCGCCTTCATGGCTCC |
| AH142 | GGAATTCCATATGGCCACCGTGGCGGAGGCG |
| KA346 | GCCAAGCTTGCGCACCGTGGCCTGACTC |
| AH143 | CCCAAGCTTGGGCAGCGGCTCCGGGCG |
| NdeI PomX fwd | GGAATTCCATATGAAGAAAGCCTTTGAACAG |
| AH144 | GCCAAGCTTTCAGGGCAGCGGCTCCGGGCG |
| KA384 | GCGGGATCCGGCGGAGCCATGAAGAAAGCCTTTGAACAG |
| DS276 | GCGAAGCTTACTTCTCGAACTGTGGGTGACTCCAGCGCACCGTGGCCTGAC |
| DS277 | GCGCCATGGCCACCGTGGCGGAGGCG |
| PomX BspHI fwd | GCGTCATGAAGAAAGCCTTTGAACAGAACG |
| PomXN rev strep-tag | GCGAAGCTTACTTCTCGAACTGTGGGTGACTCCAGGGCAGCGGCTCCGGGCG |
| NdeI-PomY fwd | GGAATTCCATATGAGCGACGAGCGTCCGGAC |
| PomY C-term his rev | CGGAAGCTTAGCGGCGAAGTATTTGTGC |
| AH141 | GCGGGATCCGGCGGAGCCGCCACCGTGGCGGAGGCG |

[1] Sequences in red indicate mutations used for site directed mutagenesis of the *pomX* gene.
